# Effective Gene Therapy for Metachromatic Leukodystrophy Achieved with Minimal Lentiviral Genomic Integrations

**DOI:** 10.1101/2024.03.14.584404

**Authors:** Lucas Tricoli, Sunetra Sase, Julia Hacker, Vi Pham, Sidney Smith, Maxwell Chappell, Laura Breda, Stephanie Hurwitz, Naoto Tanaka, Carlo Castruccio Castracani, Amaliris Guerra, Zhongqi Hou, Lars Schlotawa, Karthikeyan Radhakrishnan, Peter Kurre, Rebecca Ahrens-Nicklas, Laura Adang, Adeline Vanderver, Stefano Rivella

## Abstract

Metachromatic leukodystrophy (MLD) is a fatal lysosomal storage disease (LSD) characterized by the deficient enzymatic activity of arylsulfatase A (ARSA). Combined autologous hematopoietic stem cell transplant (HSCT) with lentiviral (LV) based gene therapy has great potential to treat MLD. However, if enzyme production is inadequate, this could result in continued loss of motor function, implying a high vector copy number (VCN) requirement for optimal enzymatic output. This may place children at increased risk for genomic toxicity due to higher VCN. We increased the expression of ARSA cDNA at single integration by generating novel LVs, optimizing ARSA expression, and enhancing safety. In addition, our vectors achieved optimal transduction in mouse and human HSC with minimal multiplicity of infection (MOI). Our top-performing vector (EA1) showed at least 4X more ARSA activity than the currently EU-approved vector and a superior ability to secrete vesicle-associated ARSA, a critical modality to transfer functional enzymes from microglia to oligodendrocytes. Three-month-old *Arsa*-KO MLD mice transplanted with *Arsa*-KO BM cells transduced with 0.6 VCN of EA1 demonstrated behavior and CNS histology matching WT mice. Our novel vector boosts efficacy while improving safety as a robust approach for treating early symptomatic MLD patients.

## INTRODUCTION

Leukodystrophies (LD) are a group of hereditary diseases affecting the central nervous system. LDs result in severe and often progressive neurologic disease, primarily affecting young children.^1,2^ The possibility of definitive, targeted gene therapies has become a reality for some LDs.^3^ One promising target of gene therapy is metachromatic leukodystrophy (MLD), an autosomal recessive lysosomal storage disease arising from biallelic pathogenic variants in the *ARSA* gene, encoding the enzyme Arylsulfatase A (ARSA), which converts 3-O-sulfogalactosylceramide (sulfatide) to galactosylceramide. Decreased ARSA activity disrupts this lipid conversion process, altering the composition of myelin sheaths and inducing a buildup of sulfatides in the CNS and peripheral nervous system (PNS).^1,4^ The most common clinical subtype is the late infantile (LI-MLD) form, which results in rapid loss of neurologic skills beginning before the age of 2.5 years and early death.^1^

Current therapy for MLD in the U.S. is HSCT^5,6^, which has mixed outcomes, especially after symptom onset.^1^ However, some European countries have approved a lentiviral (LV) treatment strategy for MLD. The drug product indicated as Atidasagene autotemcel (Libmeldy®), is made by transduction of autologous hematopoietic stem cells with an LV, indicated in this report as clinical vector (CV), encoding the ARSA gene.^7^ CV is approved for treating pre-symptomatic LI-MLD patients (onset <2.5 years) and early symptomatic patients between 2.5 and 7 years old.^8–11^ The concept behind the success of the current clinical LV is that supraphysiological expression of ARSA in pre-symptomatic patients via the Phosphoglycerate Kinase (PGK) promoter provides more enzymatic activity than allogeneic healthy hematopoietic stem cells (HSC) to suppress the detrimental effects of sulfatide buildup.^8^ A key factor in the success of this therapy is the ability of the transplanted bone marrow product to cross the BBB and replenish the endogenous microglia.^12,13^

Despite the success of the CV, there is a need to explore whether a more efficient system can be developed given the high vector copy numbers required with the currently available product. This places an increased burden on manufacturing needs and potentially a higher risk of complications such as genotoxicity for the patient. With increasing numbers of viral integration, there is a greater probability of genome toxicity and leukemogenesis, which may place children at increased risk for complications.^14,15^ Additionally, while the presymptomatic LI-MLD populations respond well to CV, treatment is inadequate to address the needs of LI-MLD early symptomatic individuals.^1^ Additional enzyme activity without increasing the need for integration may be needed to help early symptomatic patients. These limitations suggest that an improved vector design could enhance therapeutic efficacy. To this end, we generated novel vectors to be compared with the CV.

Our vector designs demonstrate greater ARSA expression and activity across biologically relevant cell lines. In addition, our vector allows generation, from the transduced cells, of secreted extracellular vesicles (EVs) loaded with the ARSA protein. *In vitro*, these EVs can transfer the ARSA protein and activity to other cells. Our LVs illustrate a robust safety profile as indicated by genotoxicity assays and transduction into WT mice. Finally, our vector ameliorates MLD disease phenotypes in *Arsa*-KO mice at a lower VCN than the CV. The combination of genetic regulatory elements that increase ARSA expression in our novel vectors can serve as a backbone for other constructs to treat similar lysosomal storage diseases and leukodystrophies, such as multiple sulfatase deficiency (MSD). Developing more efficient vectors is a critical step in reducing genome toxicity and improving outcomes for early symptomatic patients.

## RESULTS

### A novel vector design for improved transgenic expression by LVs

Several PGK and Elongation Factor 1 alpha (EF1a) promoter-driven Green Fluorescent Protein (GFP) reporter vectors were designed to assess how Ankyrin and Foamy insulators and the Woodchuck hepatitis virus posttranscriptional regulatory element (WPRE)/bovine growth hormone poly(A) signal (bGHpA, indicated as pA) sequences affected transgene expression (**Figure 1A**). The WPRE (indicated as W) is traditionally utilized to increase viral RNA integrity and titer.^16,17^ However, when included in the transgenic cassette (between the 5’ and 3’ long terminal repeats (LTR)), WPRE limits vector capacity and becomes structurally integrated into the DNA of patient cells. In addition, the Human Immunodeficiency Virus (HIV) polyadenylation signal in the 3’ LTR leaks, allowing viral RNA readthrough.^16^ As previously shown^16^, these limitations can be overcome by cloning the WPRE downstream of the 3’ LTR together with the pA. In addition, several laboratories, including ours, have documented the use of insulators to reduce position effects, silencing, and enhance safety.^18–22^ We included insulator elements in our LV design, which may minimize undesirable promoter activation and aberrant transcription in regions of LV integration.^18,19^

**Figure 1.**
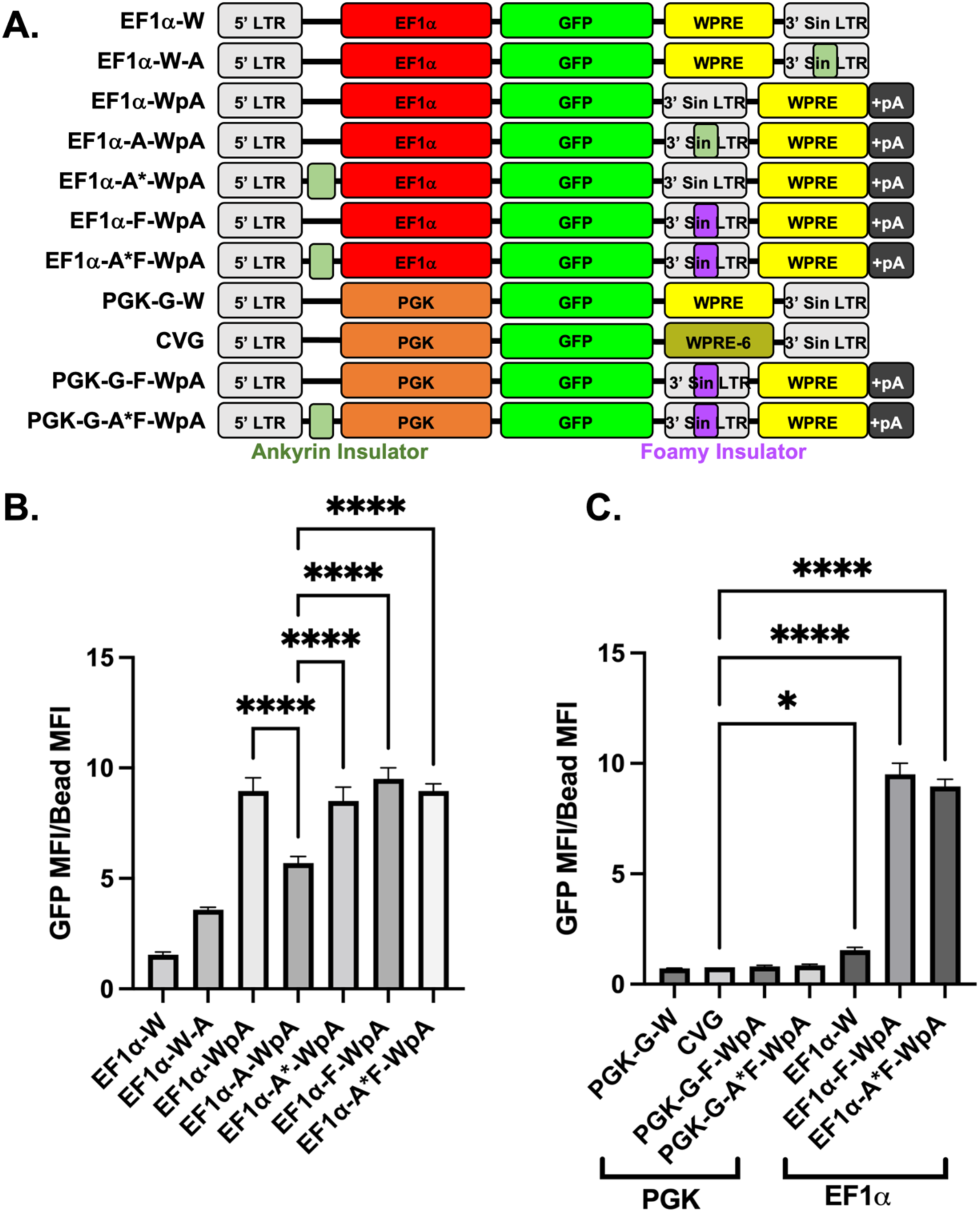
Optimal arrangement of vector components to maximize GFP expression. (A) Maps of the multiple vector arrangement used to determine the optimal configuration of EF1a promoter with ankyrin or foamy insulators and WPRE (W) with PolyA tail (pA) region for optimal GFP MFI. (B) MFI for all EF1a vector arrangements shown above (n=3). (C) MFI comparison of select PGK and the CVG arrangement compared to their EF1a counterparts to demonstrate the key expressive benefit of using Ef1a in these arrangements. LTR: long terminal repeat; Sin-LTR: self-inactivating long terminal repeat; WPRE: Woodchuck hepatitis virus posttranscriptional regulatory element; WPRE6: mutated WPRE; PGK: Phosphoglycerate Kinase promoter; EF1a: Elongation Factor 1a (EF1a) promoter; pA: bovine growth hormone poly(A) signal. *P<0.05, ****P<0.0001 Note: significance shown for key comparisons. Expanded significance is located in **Figure S1**.

In the CV, a PGK promoter drives *ARSA* expression, where a mutated WPRE (WPRE6) is downstream of ARSA and upstream of the 3’ self-inactivating (SIN) LTR.^9^ WPRE6 was generated to prevent a potential oncogenic activity of the wild type WPRE, as described in Kingsman et al.^23^ Our constructs were compared to the vector CVG, which is identical in LV format to the CV, except for replacing the *ARSA* transgene with a GFP reporter (**Figure 1A**). This allowed us to directly compare how the different genomic elements and their rearrangements enhanced transgene expression. We tested these constructs in Murine erythroid leukemia (MEL) cells, which are sensitive to integration-based position effects and transgene silencing.^18,24^

The mean fluorescence index (MFI) was measured for each vector as a metric of vector potency (**Figure 1B, 1C, S1A** and **S1B**). The percentage of transduced (positive GFP) cells was kept below 20% to avoid more than 1 integration per genome and ensure linear correlation between GFP+ cells and VCN (**Figure S2A**). Including an Ankyrin Insulator (comparing EF1a-W vs. EF1a-W-A) demonstrated a 2-fold increase in MFI (**Figure 1B**). Rearrangement of WpA (EF1a-WpA) showed an almost 10-fold increase in MFI compared to EF1a-W (**Figure 1B**). Combining insulators with WpA (EF1a-WpA vs. EF1a-F-WpA, EF1a-A*-WpA, and EF1a-A*F-WpA) had limited effects on the MFI (**Figure 1B**). Only EF1a-A-WpA showed a significant decrease in GFP synthesis (**Figure 1B**).

WpA rearrangement induced a significant increase in MFI, which correlated with a 3-fold higher GFP mRNA expression for EF1a-WpA compared to EF1a-W (**Figure S3**). Overall, the data suggests that removing the WPRE from the 3’ of the GFP gene promotes mRNA transcription and/or stability and, ultimately, GFP expression after genomic integration.

Similar modifications were introduced in PGK-based vectors (**Figure 1A** and **S2B**). None significantly impacted GFP synthesis (**Figure 1C** and **S1B**; and all PGK-driven vector MFI values and statistics are described in **Figure S2C**). However, EF1a-W demonstrated a ∼2-fold increase in MFI compared to CVG, while EF1a-F-WpA and EF1a-A*F-WpA showed almost a 10-fold increase (**Figure 1C**). This highlights the responsiveness of the EF1a promoter-driven constructs to the rearrangement and addition of LV elements compared to that of PGK. This ultimately led us to use the EF1a-A*F-WpA arrangement of elements to design a more potent expressing *ARSA* vector compared to the construct of the CV.

### Novel LV vectors increase *ARSA* expression and activity in patient-derived fibroblasts and *ARSA*-KO human microglia cells

Based on the preliminary results with the GFP reporter constructs, we generated six new vectors to drive *ARSA* expression and compared them to an in-house replica of the approved clinical vector for MLD (cloned according to the published literature on Atidasagene autotemcel), indicated as CV (**Figure 2A**).^11^ The ARSA coding sequence was optimized to increase its translation in human cells and/or generate a coding sequence that could discriminate between patient and vector *ARSA* mRNA by RT-PCR (E1-E4). Additional modifications were applied to generate EA1 and EA2, which contained the foamy insulator in the 3’ LTR with and without the ankyrin insulator 5’ of the EF1a promoter. Both vectors contained the sequence-modified ARSA (without UTRs) to retain the ability to discriminate between endogenous and transgenic mRNA sequences, described in **Figure 2A**.

**Figure 2.**
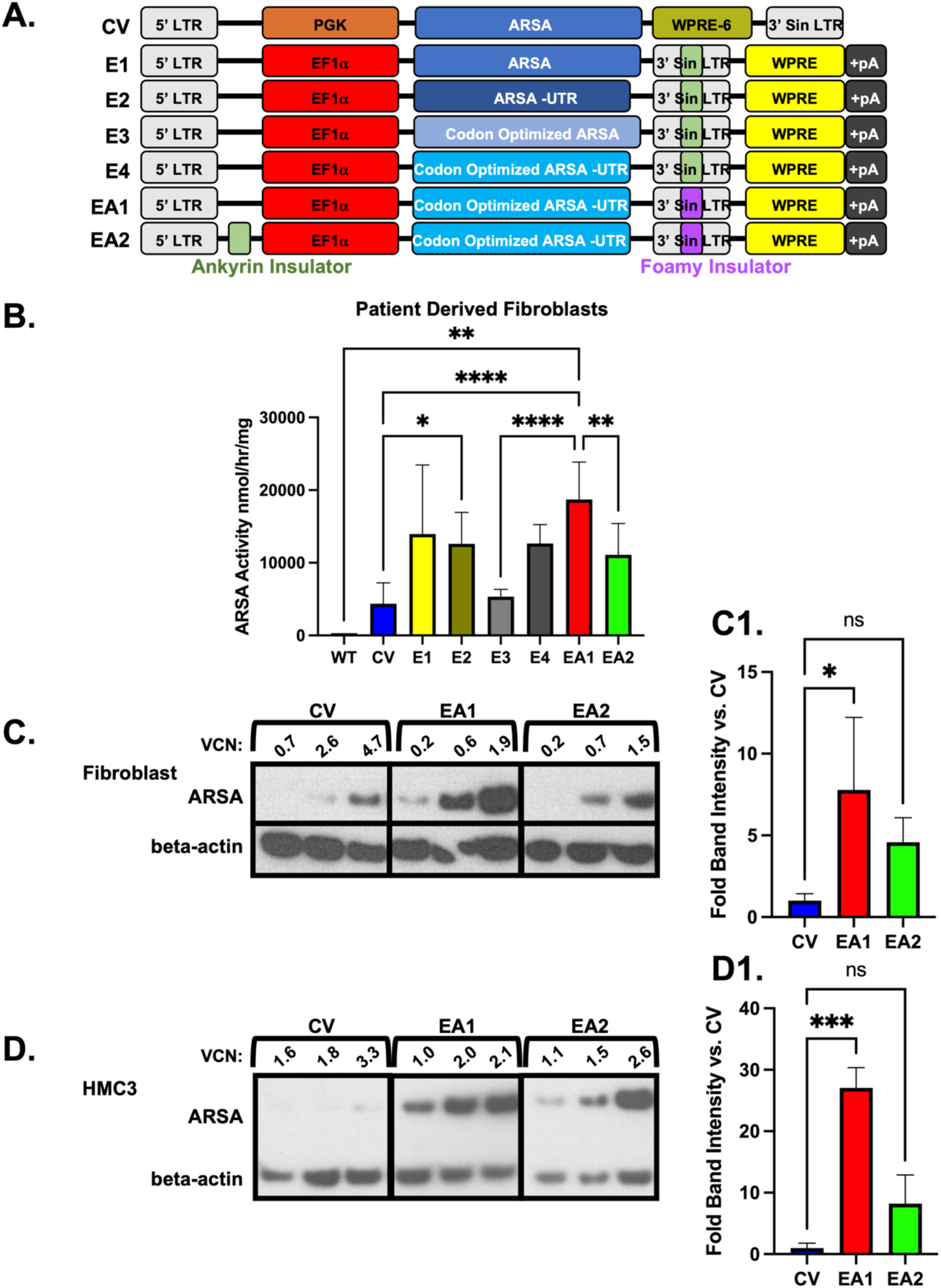
Improved activity and protein expression of ARSA in MLD patient-derived fibroblasts and mouse HSCs. (A) Vector schematics comparing the CV and our six Ef1a vectors used to test ARSA expression based on the best MFIs from the GFP pilot vectors. (B) Fibroblast ARSA enzyme activity normalized to VCN for our 6 test vectors, including the CV (n = ranged from 3-10). (C) A western blot of ARSA expression from the same transduced patient-derived fibroblast in (B). (D) A western blot of ARSA expression run for a human ARSA KO microglia cell line (HMC3) transduced with CV, EA1 and EA2 vector. (Fold change in band intensity is normalized to beta-actin). (E) Quantification of ARSA expression in fibroblast and HMC3 by western blots shown in (C) and (D) (n = 3). LTR: long terminal repeat; Sin-LTR: self-inactivating long terminal repeat; WPRE: Woodchuck hepatitis virus posttranscriptional regulatory element; PGK: Phosphoglycerate Kinase promoter; EF1a: Elongation Factor 1 alpha (EF1a) promoter. *P<0.05, **P<0.01, ****P<0.0001

WT and *ARSA*-deficient primary fibroblasts (cell line GM02331) derived from a healthy individual or a patient with MLD, respectively, were used to test ARSA expression and enzymatic activity upon transduction with either CV or one of our vectors. We analyzed ARSA activity in specimens with transduction of VCN 2 or lower to evaluate the correlation between ARSA activity and VCN as a linear relationship. CV-treated cells demonstrated enhancement in ARSA activity over WT cells (**Figure 2B**), concordantly with data previously published.^8,9^ Comparing E1, E2, E3, and E4 vectors that contained the ankyrin insulator, neither optimization of the coding sequence nor inclusion of the 5’ and 3’ UTRs significantly modified ARSA synthesis. In addition, although we observed a trend in increased ARSA activity in most of these vectors compared to CV, the differences were not significant (except for E2) (**Figure 2B**). EA1-treated cells showed 4-fold greater activity in ARSA-deficient fibroblasts than those treated with CV (**Figure 2B**). Therefore, EA1 was chosen for further studies. EA2 was also selected because of its structural similarity to EA1 and to test the effect of expressing an intermediate level of ARSA (compared to EA1 and CV). EA1- and EA2-treated fibroblasts demonstrated more transgenic ARSA expression than those treated with CV (**Figure 2C**). EA1 and EA2 treated specimens synthesized, on average, 7-fold and 4-fold greater ARSA protein (normalized to beta-actin and VCN, expanded description in the materials and methods) than specimens treated with CV (**Figure 2C** and **2C1**). EA1 was assessed in transduced human CD34+ cells (**Figure S4**), showing at 1.8 VCN ∼3x more ARSA expression than CD34+ cells alone.

Transfer of ARSA protein to MLD-affected cells in the central nervous system (CNS) by HSC-derived microglia may contribute to treatment efficacy through cross-correction but has not definitely been proven to play a role.^25,26^ Therefore, ARSA synthesis and activity were also assessed in the *ARSA* knock-out (KO) human microglia cell line (HMC3) exposed to EA1, EA2, or CV. As shown in **Figures 2D** and **2D1**, ARSA expression of EA1 and EA2 treated specimens was, on average, 27-fold and 8-fold greater (again normalized to beta-actin and VCN) than that measured in CV-treated cells (1.8 VCN), further confirming superior ARSA expression over CV.

### EV secretion as a modality of ARSA enzyme transfer in the CNS and superior ARSA production by EA1 in EVs

Secreted small extracellular vesicles (EVs) ranging from 50-200 nm in size have been well demonstrated to transfer functional biological cargo from host to recipient cells. Clinical interest has thus emerged in engineering EVs for therapeutic delivery of functional enzymes in lysosomal storage disorders, though the role of cross correction in treatment remains unknown.^26,27^ Cellular overexpression of target cargo may facilitate physiologic loading and secretion of constitutive EVs from modified cells.

We evaluated secretory levels of ARSA from cell-conditioned media of primary patient MLD fibroblasts and *ARSA*-KO microglia after transduction with CV or EA1. EA1-transduced cells demonstrated an increase of EV-secreted ARSA at comparable VCN to CV-transduced cells (**Figure 3A**). Cell-free conditioned media was depleted of beta-actin and enriched in CD81 levels, a tetraspanin protein associated with secreted EVs.

**Figure 3.**
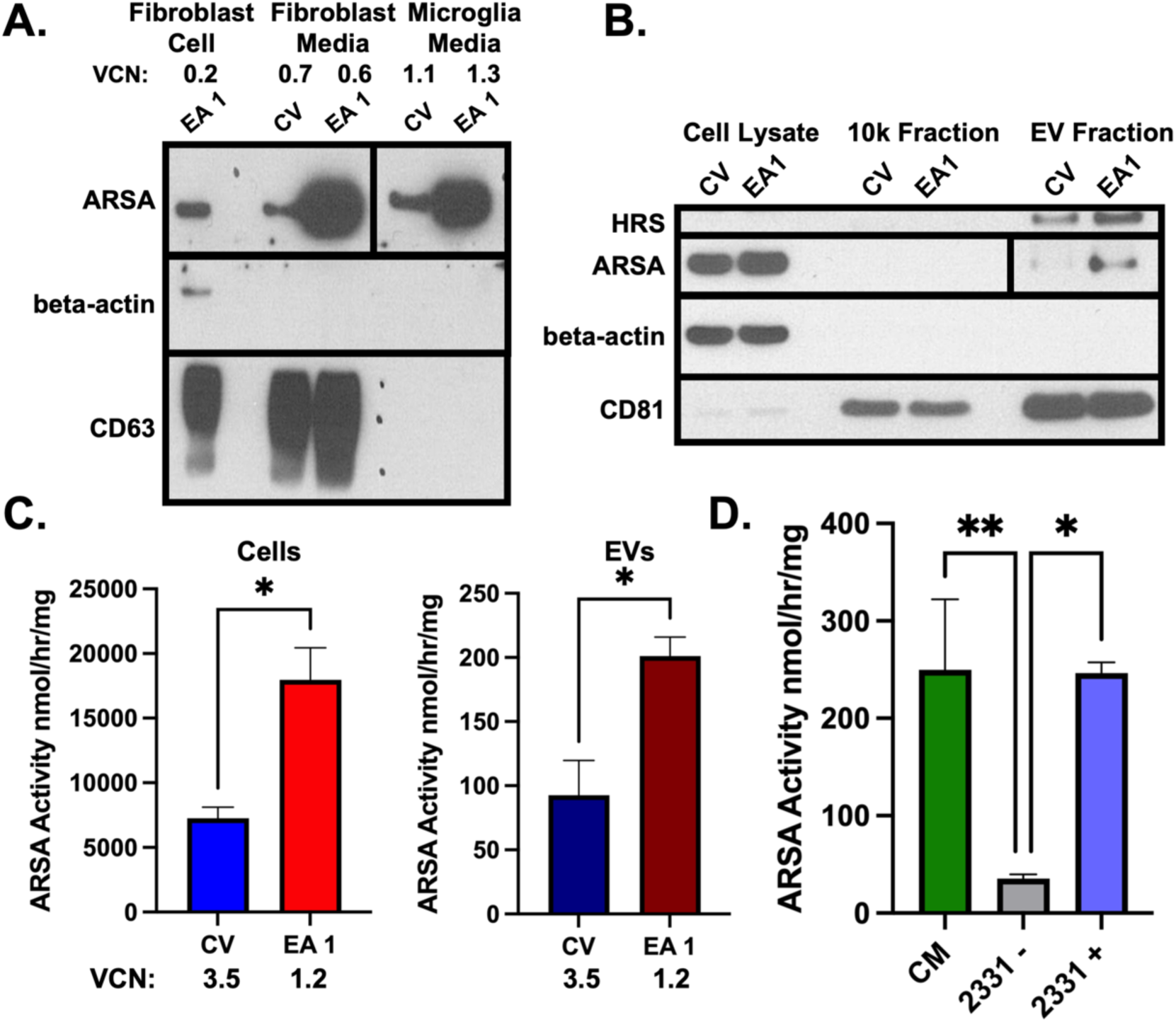
Increased ARSA protein expression and activity in EVs secreted from transduced cell lines. (A) Protein-precipitated media harvested from vector-transduced patient fibroblasts and *ARSA*-KO microglia cells demonstrated much greater ARSA expression in EA1 than CV by western blot. The VCN of each cell line is noted above the treatment (top left). (B) EVs were then isolated via ultracentrifugation, yielding more expression of EV-related ARSA for EA1 (VCN 1.2) compared to CV (VCN 3.5). The cells the media was derived from (cell lysate) were also shown for comparison in addition to the 10k xg large vesicle fraction. (C) ARSA activity assay for the same cells demonstrated higher ARSA activity in cells per VCN as demonstrated previously and more ARSA activity from secreted EVs (n = 2). (D) ARSA activity of media conditioned by an EA1 transduced patient fibroblast line (CM). ARSA activity of the untransduced patient cell line alone (2331-) compared to when CM is added to the untransduced cells (2331+), an increase in ARSA activity to the transferred cell line is observed (n = range 2-5). *P<0.05

To measure vesicle-associated ARSA abundance and activity, secreted EVs were isolated from human MLD primary fibroblasts transduced with EA1, EA2, or CV. Levels of ARSA were mainly enriched in small, secreted hepatocyte growth factor-regulated tyrosine kinase substrate positive (HRS) and CD81^hi^ (both markers of EV enrichment) vesicles isolated after ultracentrifugation (100K xg) compared to larger HRS-CD81^lo^ vesicles isolated after a low-speed (10K xg) centrifugation (**Figure 3B** and **Figure S5A**). Density gradient-purified small EVs from EA1 and CV-transduced cells were enriched in ARSA, with detectable levels in EA2-transduced EVs (**Figure S5B**).^28,29^ EA1-transduced cells showed an increased amount of secreted enzyme at lower VCN (EA1:1.2 versus CV:3.5). Functional ARSA activity was enhanced about 2-fold in EA1-transduced cells and secreted vesicles compared to CV-treated cells (**Figure 3C**). Conditioned media from EA1-transduced cells was added to *ARSA*-deficient patient fibroblasts to test whether EV-associated ARSA could be functionally transferred. This yielded significant increases in cellular enzyme activity (**Figure 3D**). These data indicate that EA1 transduction enhances EV-associated secretion of functional ARSA compared to CV. This increased efficiency of the secretory pathway will likely impact the enzymatic load delivered to non-hematopoietic cells, which may improve outcomes after gene therapy.

### EA1 and EA2 vectors exhibit no observable immortalization in transduced HSCs

In addition to maximizing ARSA expression and activity, vector safety is a vital feature in preclinical consideration. To exclude genome toxicity *in vitro*, we evaluated our vectors’ potential to transform lineage-negative selected mouse bone marrow cells in long-term replating experiments, as shown previously.^30^

Lineage-negative selected mouse HSCs were transduced with CV, EA1, and EA2. LV transduced mouse HSC cells were compared to mock un-transduced cells, and as a positive control, we used a well-established gamma retroviral vector named pMSCV-GFP.^30,31^ We did not observe clonal expansion in any of the lentiviral transduced cells, up to 6.4 VCN, while as expected, pMSCV-GFP induced clonal expansion starting at week 5 and VCN starting at 1.7 **(Figures 4A** and **4B)**. For the pMSCV-GFP vector, the transformation frequency was proportional to the VCN (**Figure 4B**).

**Figure 4.**
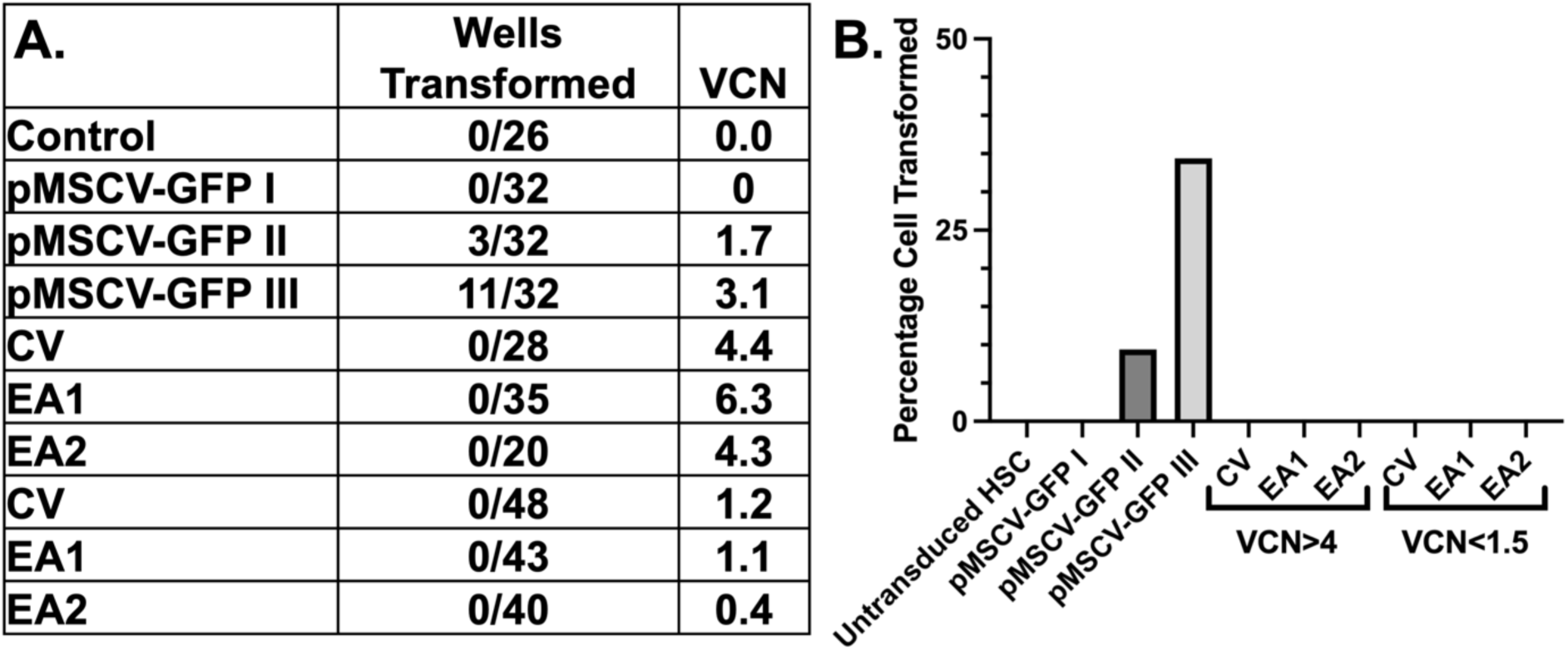
Genotoxicity assay. (A) A table of VCN and percentage of wells transformed in multiwell well plate format of vector transduced mouse HSCs. The positive control pMSCV-GFP vector-treated wells are the only plates with cell transformation and media depletion (yellowing) due to cell immortalization. All other vector-transduced plates after 9 weeks show cellular senescence and absence of media depletion. (B) A graphical representation of absence in cell transformation in all cell lines aside from the pMSCV-GFP, indicating the safety of our vector at higher (>4) and clinically relevant (1.5>) VCN.

### EA1 and EA2 vectors do not perturb physiological hematopoiesis in transplanted WT mice

We further explored the safety of our vector by transplanting WT mice with GFP donor marrow alone or transduced with EA1, EA2, and a modified CV vector, where ARSA was replaced by GFP (CVG from **Figure 1A**).

Several animals infused with GFP mouse bone marrow transduced with EA1 and EA2 showed high VCNs (VCN>5; **Figure 5A**) analyzed approximately 5 months post HSCT.

**Figure 5.**
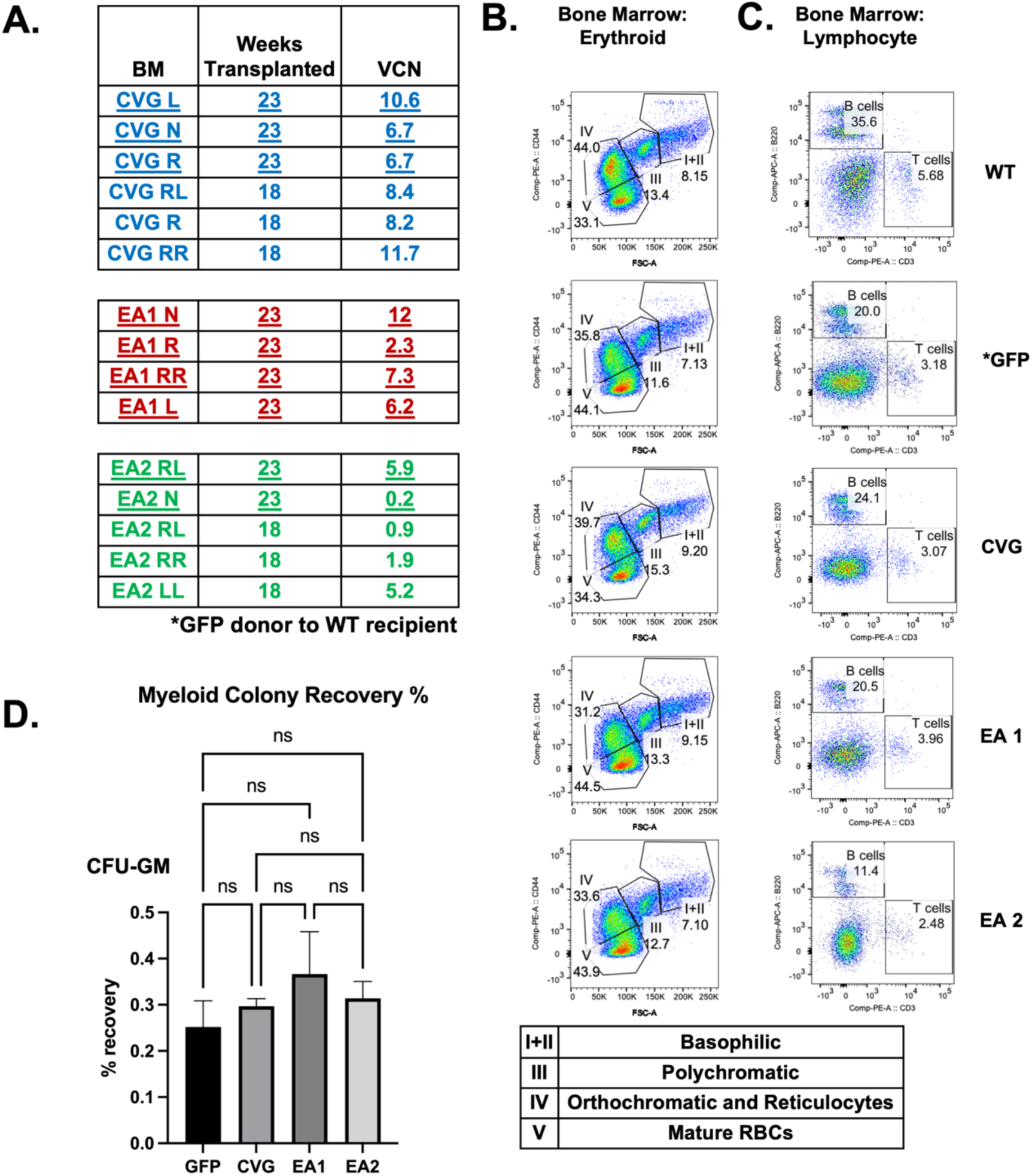
Transduction of GFP CV, EA1, and EA2 vectors into WT mice demonstrate vector safety. (A) The number of weeks post-BMT and VCNs are indicated. Mice chosen for the CFU assay are underlined. (B) Example erythropoiesis analysis (population I to V, from immature to enucleated red cells) by flow cytometry in WT controls, GFP mice transduced with normal marrow or mice treated with CVG, EA1, or EA2, from top to bottom. (C) In the same animals, the lymphocyte B and T cell immunoprofiles are also shown. (D) Myeloid progenitor colony formation assay from transduced GFP mouse mock control, CVG, EA1, and EA2 treated mice. CFU-GM = granulocyte-macrophage progenitor.

We did not observe gross abnormalities in erythroid or lymphocyte populations between animals engrafted with transduced or untransduced HSC and WT control mice (**Figure 5B**). The percent distribution of erythroid cells at different stages of maturation (**Figure 5B**) was similar among cohorts. Also, lymphoid B and T cell populations demonstrated a comparable percentage distribution in vector-transduced mice compared to controls (**Figure 5C**, with additional flow profiles for every harvested mouse described in **Figure S6A** and **S6B**). A mouse bone marrow colony forming unit (CFU) assay also showed no difference between treated and control mice (**Figure 5D**).

### *Arsa*-KO mouse disease model of MLD is corrected with, on average, 5-fold less EA1 vector than CV

We generated a novel *Arsa*-KO mouse model of MLD, as described in **Figure S7**. This mouse model is expected to recapitulate the phenotype of a previously generated mouse model, which was not available for our studies.^32^ In pilot experiments, after eight months, we observed significant sulfatide accumulation and microglia inflammation in *Arsa*-KO brains, hallmarks of MLD pathology (**Figure S8A**). However, at 8 months *Arsa*-KO mice did not demonstrate a significant difference in rotarod performance compared to a control mouse. The motor deficit became apparent in one-year-old *Arsa*-KO mice, which showed a significant difference on the rotarod, corroborating evidence of disease phenotype and the need to assess mice at one year of age **(Figure S8B**).

With these parameters in mind, we compared the efficacy of therapeutic vectors: 8- to 12-week-old *Arsa*-KO lethally irradiated mice were infused with *Arsa*-KO HSC transduced with CV, EA1, or EA2 or control *Arsa*-KO HSC. Untransduced *Arsa*-KO and WT were used to identify baseline values. Phenotypic rescue was evaluated ten months post-HSCT, at approximately 12 months total age.

Our study aimed to design a vector to prevent the MLD disease at lower VCN. Therefore, challenging our vector to outperform CV at lower VCN was critical. Given the potency of EA1 observed *in vitro*, we aimed for a VCN of around 1. On rotarod testing, WT mice showed improvement in their maximum duration over the course of the 5-day testing period, starting at about 230 seconds and going to almost 260 by day 5 (**Figure 6A**). In comparison, the *Arsa*-KO mice duration started the testing period with reduced maximum duration at baseline compared to WT and demonstrated some improvement over the testing period by day 5 (**Figure 6A**). CV-treated mice began at 200 seconds in duration and improved very little by day 5, generally performing poorly compared to WT (**Figure 6A**, **Table S1**, **Figure S9A** and **S9B**). EA1 and CV-treated mice were split into 2 VCN groups (**Figure S9**, **Tables S1**, and **S2**), highlighting a bifurcation in mouse phenotypic improvement based on the VCN administered. At a VCN range of 1.1-2.6, CV-treated mice performed similarly to WT mice on the rotarod. It is important to note that the 2 VCN range is the clinically relevant VCN for the efficacy of this vector.^7^ In contrast, the performance of EA1-treated mice matched that of the WT group, at an average VCN of 0.35 (**Figure 6A**), about 5-fold less than that of the 1.1-2.6 VCN CV-treated mice. When plotting the duration time on the rotarod for day 5 against VCN, we found a linear correlation between these parameters in the CV cohort, which was confirmed by the slope being significantly different from zero by statistical linear regression analysis (**Figure 6B**).

**Figure 6.**
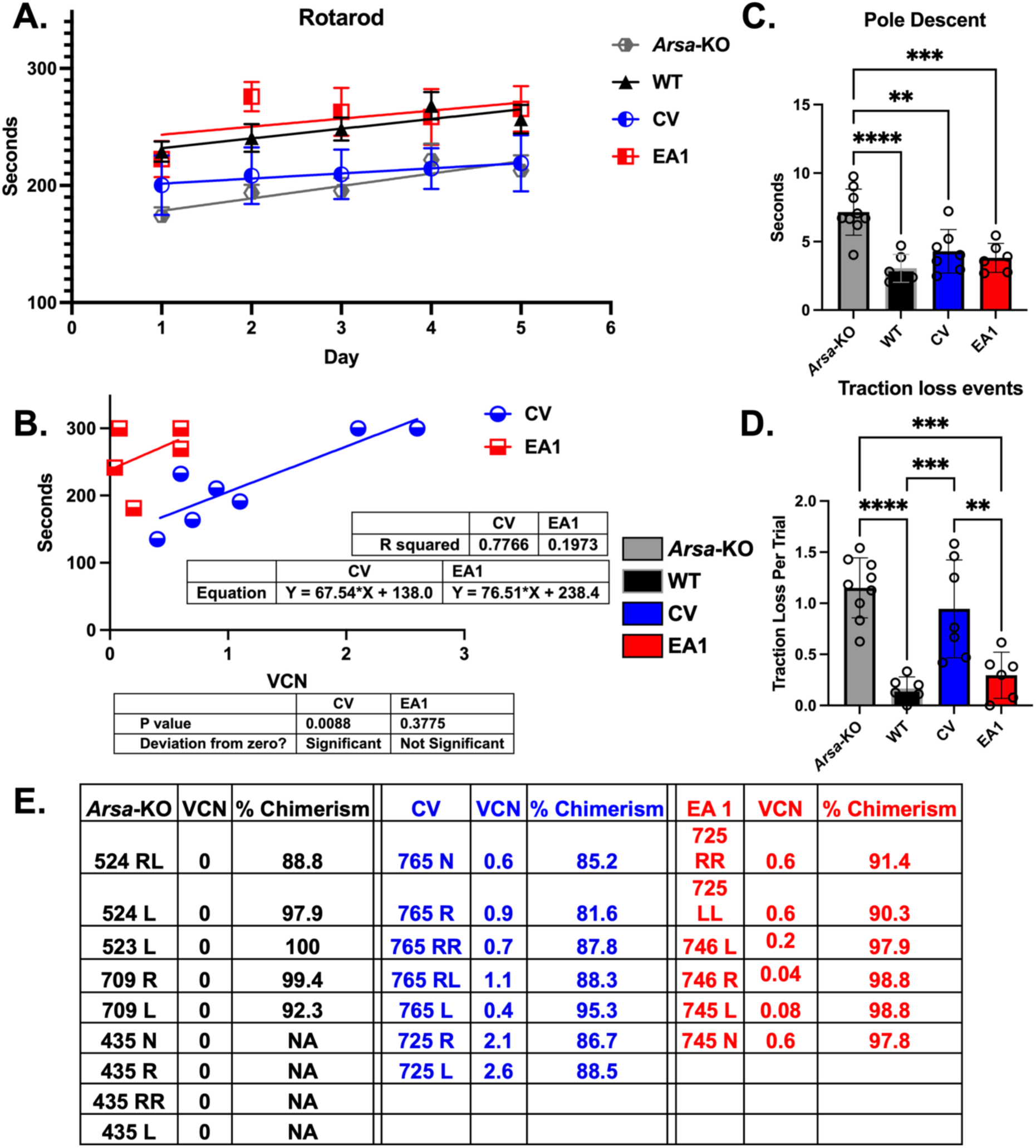
Rotarod and pole descent motor control assays of untreated *Arsa*-KO disease mice compared to vector-treated mice. (A) Rotarod assessment over a 5-day time course for WT, untreated, or CV or EA1 vector treated mice (n = range 6-9). (B) A linear correlation demonstrating the relation between VCN and mouse duration (performance) on the rotatord at day 5 of assessment. (C) Descent time duration for a predetermined pole segment in vector-treated groups compared to *Arsa*-KO mice (n = range 6-9). (D) Report of times mice slipped or lost grip during the pole descent assay (n = range 6-9). (E) Table of mice assessed and corresponding VCN and chimerism. *P<0.05, **P<0.01, ***P<0.001, ****P<0.0001. Extended rotarod significance tables in supplement (**Tables S1** and **S2**).

We did not observe a linear relationship between rotarod duration and VCN for EA1-treated mice.

As a baseline in the pole descent assay, WT mice demonstrated significantly more rapid descent times than *Arsa*-KO mice, traversing the pole in about 3 seconds on average (**Figure 6C)**. WT mice performed with less than half of traction loss event per trial during descent compared to *Arsa*-KO mice (**Figure 6D**). In comparison, *Arsa*-KO mice demonstrated the longest descent times, averaging around 7 seconds, and significant difficulty gripping the pole, with an average of over one traction loss event per trial during the descent (**Figure 6C** and **6D**). CV-treated *Arsa*-KO mice had an average descent time of over 4 seconds and significantly differed from untreated *Arsa*-KO mice. Data was bifurcated into cohorts CV 0.4-0.9 VCN, CV 1.1-2.6 VCN, EA1 0.04-0.2 VCN and EA1 0.6 VCN in **Figures S9C** and **S9D**. The CV cohort with higher VCN did demonstrate the biggest improvement in descent times in line with WT mice, while the CV cohort at lower VCN showed no differences compared to *Arsa*-KO. However, the overall traction loss events for all CV-treated cohorts were similar, showing no significant difference to *Arsa*-KO mice, again highlighting a lack of complete disease prevention for CV irrespective of VCN. The EA1 cohorts showed more significance than CV compared to *Arsa*-KO for descent times, with an average descent of 3.8 seconds. Traction control on the pole during the EA1 group’s descent was in line with WT mice. The marked improvement in motor function with EA1 treatment was accomplished with an average of only 0.35 VCN of vector integration across all VCN-treated groups compared to 2 VCN for CV cohort with the high VCN. These data consolidated EA1 as our best vector and highlighted the 5-fold increase in vector potency, which aligns with the *in vitro* results. The chimerism in transplanted mice was above 80% and comparable among experimental groups (**Figure 6E**). Secondary transplants were performed from primary transduced EA1 mouse whole BM, and no gross abnormalities in hematopoiesis were demonstrated by flow cytometry (**Figure S10**).

### Histopathology demonstrates correction of MLD phenotype in line with behavioral analysis

To determine if lower VCN in treated EA1 mice with improved behavior is accompanied by amelioration of brain physiology, we performed neuropathology on the brain sections of the same treated mice at 12 months of age. *Arsa*-KO mice presented many sulfatide bodies in the corpus callosum and other brain areas relative to WT mice (**Figure 7A**). When plotting the correlation of sulfatide body accumulation to VCN, we found a similar trend as we did in the rotarod behavior data in that mice treated with CV presented a reduction of sulfatide bodies at a higher copy number than that of EA1-treated mice (**Figure 7A1**). Quantification of sulfatide bodies indicated that EA1-treated mice dosed with lower VCN (at 0.6) had counts in line with WT mice. In the CV cohort with lower VCN (0.6) sulfatide bodies demonstrated more accumulation compared to that of the EA1 cohort (**Figure 7A** and **7A2**). The CV cohort with higher VCN showed no sulfatide bodies and efficiently corrected this phenotype (**Figure S11A**).

**Figure 7.**
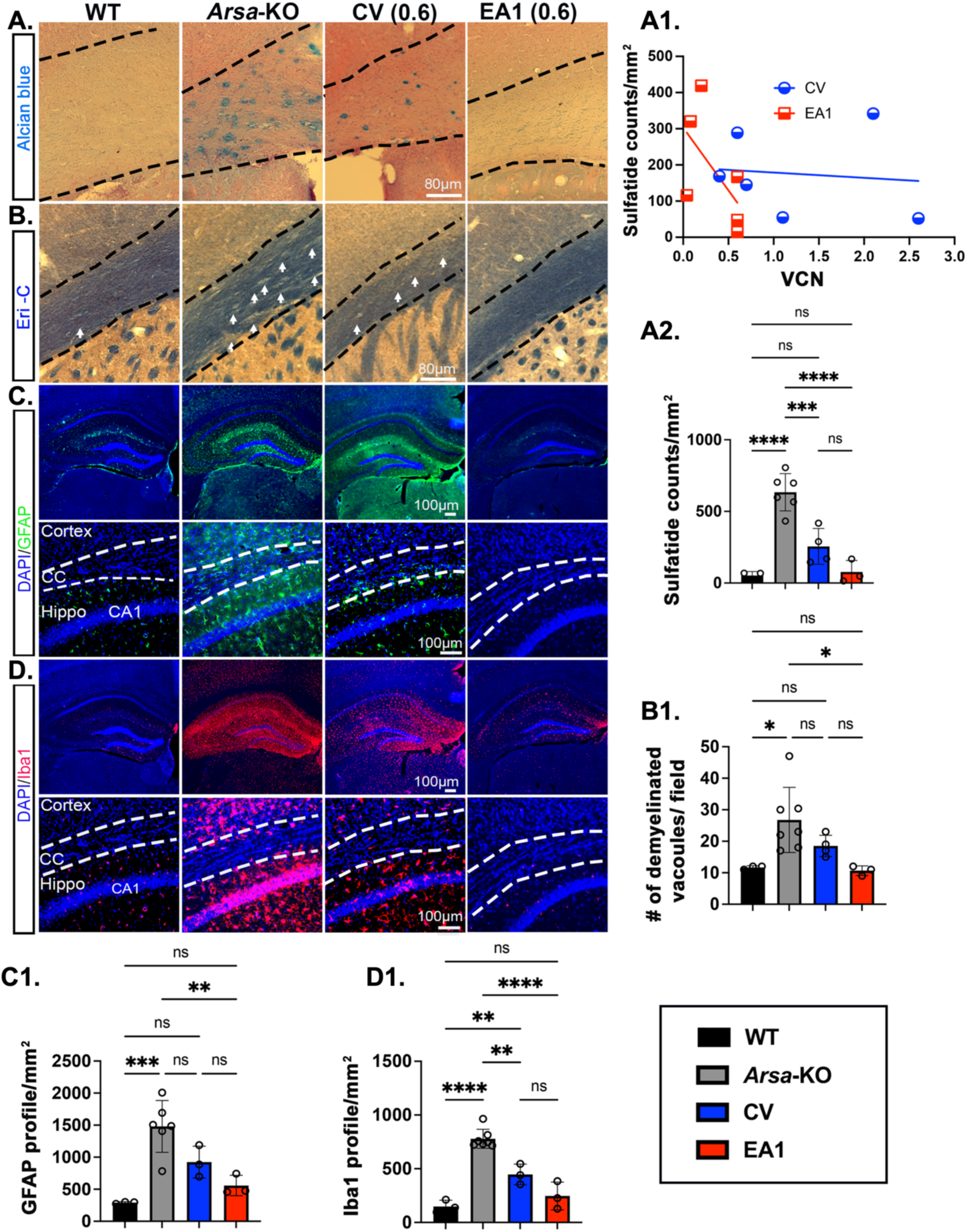
Neuropathological deficits of untreated *Arsa*-KO diseased mice relative to vector treated mice. Treatment groups include WT, untreated, CV, and EA1 vector treated at lower VCN. (A) Representative images of sulfatide accumulation in the corpus callosum across treatment groups. (A1) Linear correlation exhibiting the relation of VCN and sulfatide accumulation in the corpus callosum of CV and EA1. (A2) Quantification of sulfatide accumulation across groups. (B) Representative myelin staining (Eri-C) images of the corpus callosum in treatment groups. (B1) Graphical presentation of # of demyelinated vacuoles (as shown by white arrows in (B)). (C) Representative immunofluorescent images of GFAP (astrocytes) in different brain areas. (C1) Astrocytes (GFAP) counts/mm^2^ in the corpus callosum. (D) Representative immunofluorescent images of Iba1 (microglia) in different brain areas. (D1) Microglia (Iba1) counts/ mm^2^ in the corpus callosum (n=3-6 per group). *P<0.05, **P<0.01, ***P<0.001, ****P<0.0001. Extended significant tables in **Table S3**.

Next, we evaluated myelin deficits and patterns using Eriochrome cyanine (Eri-C) staining in *Arsa*-KO mice. *Arsa*-KO mice demonstrated demyelination spots or vacuoles (**Figure 7B**, indicated by white arrows) compared to WT mice. We quantified these demyelination vacuoles in untreated and treated groups (**Figure 7B1**). Mice treated with EA1 at VCN of 0.6 showed no vacuoles and well-preserved myelin. In comparison, mice treated with CV at lower VCN failed to show complete correction in myelin. Only at VCN > 1.1 CV-treated mice showed improving phenotypic features (**Figure 7B1, Figure S11B,** and **Table S3B**).

Further, to evaluate neuroinflammation in *Arsa*-KO mice, we performed GFAP and Iba1 staining in astrocytes and microglia. *Arsa*-KO mice showed increased astrogliosis and microgliosis relative to WT mice (**Figure 7C** and **7D**). In the treated groups, EA1-treated mice at lower VCN (0.6) exhibited no neuroinflammation in different areas; instead, CV-treated mice with equivalent VCNs demonstrated some astrogliosis and microgliosis (**Figure 7C1** and **7D1**). At higher VCNs, CV-treated mice demonstrated complete correction (Figure **S11C** and **S11D**).

Overall, our novel EA1 vector is highly efficacious in ameliorating the disease phenotype at considerably low VCN.

## DISCUSSION

We have engineered a novel LV for autologous hemopoietic stem cell gene therapy of MLD that provides more robust ARSA expression at lower VCN compared to a copy of the clinical vector currently approved in the EU for MLD, here named CV. *In vitro*, the EA1 vector produces 4x more ARSA activity per VCN than CV. This prevented the development of the MLD phenotype in *Arsa*-KO mice with, on average, 5x fewer copies per genome (VCN 0.35) of EA1 than CV (VCN 2). In previous experiments with a vector like CV, a protective effect in *Arsa*-KO mice was achieved with VCN in the range of 3-9.^8^ In patients, a vector-like CV produces a protective effect closer to the 2 VCN range.^7,9^ This highlights the need for a more robust vector to accomplish the same effect at a much lower VCN. The low VCN in MLD animals rescued by EA1 could limit possible genotoxic effects and improve manufacturing efficiencies. There is also the potential that a more efficient vector could require less myeloablation, further reducing potential therapy-related complications. With the Adrenoleukodystrophy lentiviral product, the risk of myelodysplastic syndrome and trial suspension highlight the need to develop safer gene therapy vectors.^33^ We have taken further precautions by including insulator elements to help mitigate insertional genotoxicity.^20,21^ Achieving the most expression possible while maintaining safety should be the benchmark for new cutting-edge gene therapies.

In our hands, EA1 transduces human CD34+ cells very efficiently, with a minimal MOI of 10, which yielded a VCN of 1.8. In transplanted mice, EA1 VCNs in the range of 1 were achieved with ∼0.5 MOI. Given the current hurdles and costs in LV production, our vector provides a strong case for being less expensive (reduced MOI to achieve optimal transduction), and efficient (low VCN to be effective).^34^ Compared to CV, if a lesser amount of vector can be implemented to achieve the same effect, this could drastically reduce the cost of treatment. Atidasagene autotemcel is one of the most expensive therapies, at about 3 million EUR per treatment.^35^ This can pose a substantial financial burden to families receiving the treatment. Given that LV treatments are becoming increasingly mainstream as a highly effective treatment strategy, we must strongly consider making our vectors more competitive for widespread and affordable implementation.^36^ The amount of CV integration required for effective treatment is a critical consideration for viable treatment of MLD. For the first time, we have demonstrated that the CV has a limited effective range in protective ability to prevent MLD in an *Arsa*-KO mouse model. Mice treated at around and below 1 (at low) VCN showed no amelioration in behavior on rotarod analysis, pole descent, and traction loss events and performed similarly to the untreated *Arsa*-KO mice overall.

Cross-correction and transfer of functional ARSA enzyme from microglia to the surrounding CNS is a potential treatment modality, although this is controversial.^25,26^ We determined EVs to be a significant source of ARSA secretion. In vitro, studies of conditioned media containing EVs demonstrated a modality for ARSA enzyme transfer from transduced cells to untransduced ones. EA1 produced more functional ARSA per VCN in EVs than CV. Microglia repopulation into the CNS from the BM is a crucial feature of MLD treatment.^12^ The CNS niche created after myeloablation must be populated with microglia that produce optimal ARSA to mitigate the disease’s onset.^12^ Since there is only one opportunity for the adoptive transfer of autologous gene-modified cells, the amount of ARSA secreted from microglia should be maximized for delivery into other CNS cells.

As we advance therapy development across the leukodystrophies (and rare diseases in general), we face serious limitations regarding vector production and concerns about increased risk with increased copy requirements. We propose that a more efficient vector system, such as EA1, may meet the growing needs of the rare disease community. This has the potential to also support populations that are not currently served by commercial vectors, including symptomatic and late-onset populations. Currently, the existing Atidasagene autotemcel is approved only for pre-symptomatic patients. This, in turn, requires neonatal/prenatal screening or a familial history of MLD to address this population of children. However, widespread identification of the broad panel of rare diseases, such as MLD and other lysosomal storage disorders, is still lagging.^37,38^ Therefore, being adequately prepared to overcome diseases at any stage is critical for successful treatment. With our more potent EA1 vector, we may be able to offer options for children of families currently with no alternative and address a population over a much broader range of clinical manifestations.

The effectiveness of the improved backbone and regulatory elements in LV design^1,37,38^ has a potential impact on other related leukodystrophies like MSD, as described in the manuscript by Vi Pham and colleagues (companion paper). The ability to address multiple associated diseases with a single vector backbone highlights the versatility and advancement of our lentiviral vector program.

## MATERIALS AND METHODS

### Cell Lines

MLD patient fibroblast cell line, designated GM02331, was generated by Adeline Vanderver (CHOP) and maintained in MEM supplemented with 15% non-heat inactivated FBS with 10% pen strep. *ARSA*-KO microglia cell lines were generated using human HMC3 cells. CRISPR was utilized by Synthego to develop *ARSA*-KO cells as according to the strategy outline in **Table S4**. Cells were then cultured according to ATCC guidelines, and single cells were sorted to derive a homogeneous population of characterized HMC3 *ARSA*-KO cells. Colonies were expanded and sequenced to verify the KO of ARSA and the lack of ARSA synthesis. One of these colonies was selected as our model microglia *ARSA*-KO line and indicated as HMC3 14.

### Plasmid, virus production, and transduction of MEL cells

We used PGK or EF1a promoters to drive transcription of the gene encoding green fluorescent protein (GFP) to assess more robust vector constructs. Vectors were constructed with a GFP reporter to determine the Mean Fluorescence Index (MFI).^19^ LV constructs were designed in our laboratory, and all plasmids were manufactured by Genscript (Piscataway, NJ). Plasmid DNAs were expanded and isolated using a Qiagen Maxiprep kit (Cat 12162, Germantown, MD).

Our LV vectors were assembled using a 3^rd^ generation LV system (VSVG, REV, and RRE) in 293T cells and concentrated via ultracentrifugation.^16^ GFP vector construct transductions were conducted in MEL cells.^18^ Murine erythroleukemia cells were grown in RPMI (Cellgro, Corning, Glendale, Arizona) and supplemented with 10% FBS (Hyclone, South Logan, UT) and 1% penicillin/streptomycin. Transduction was conducted at 5x10^5^ cells/mL concentration, complemented with 1% polybrene (Millipore, Billerica, MA).

MEL cell erythroid differentiation was achieved by seeding 2×10^6^ cells, in log phase, into 3 mL of fresh culture media containing HMBA at a final concentration of 1 mg/mL. Treatment with N,N’-hexamethylene bis(acetamide) (Millipore Sigma, Burlington, MA) (HMBA) was repeated after 2 and 4 days of culture, with cells being collected for analysis after seven days. Mel cells were transduced with a low titer of virus and were assessed to have under 20% GFP% by flow (**Figure S2**). This was essential to ensure MFI was proportional to one integration per cell (under 20% GFP +). BD FACSCanto (Piscataway, NJ) was used for flow cytometry analysis.

### ARSA Activity Assay

ARSA activity was assessed through a nitrocatechol colorimetric sulfatide assay.^39^ ARSA activity was measured in nmol/hr/mg. ARSA colorimetric activity assay was conducted as outlined by Baum et al.^39^ Assay details were taught by David Wenger from Thomas Jefferson University and adopted to a 6-well plate by us. Briefly, frozen pellets were sonicated at 50% amplitude in one-second pulse on and one-second off for six pulses. 4-Nitrocatechol substrate, phosphate reaction buffers, and NaOH solutions were all prepared as outlined by Baum et al.^39^ After reaction quenching, 300 uL of the liberated nitrocatechol reaction volume was added to a well of a 96-well plate, and absorbance was read at 515 nm.

### Western blot analysis

Protein was quantified with Pierce BCA Protein Assay Kit (Thermo Fischer Scientific, Waltham, MA). Gels were run on a NuPAGE 4-12% Bis-Tris 1.5mm x 12 well gel (Thermo Fischer Scientific, Waltham, MA). The transfer was conducted onto a PVDF membrane (Bio-Rad, Hercules, California). Gels were probed with antibodies against hARSA (R&D systems, Minneapolis, MN), CD63 (Abcam, Waltham, MA) beta-actin (Santa Cruz, Dallas, TX), HRS (Santa Cruz, Dallas, TX), Calnexin (Santa Cruz, Dallas, TX), Rab5 (Abcam, Waltham, MA), Syntenin-1 (Abcam, Waltham, MA) CD81 (Santa Cruz, Dallas, TX), and CD9 (Millipore, Darmstadt, Germany), then visualized with SuperSignal Extended Duration Substrate (Thermo Fischer Scientific, Waltham, MA). Audioradiography films (Thomas Scientific, 1141J52, Swedesboro, NJ) band densitometry were analyzed by ImageJ (https://imagej.net/ij/index.html). Beta-actin band intensities were normalized internally to the lowest beta-actin signal for each vector treated sample VCN series. Human ARSA band intensity were then normalized to beta-actin for each lane. Finally, each beta-actin normalized ARSA band intensity was normalized to its respective VCN. These normalized values were then used to determine the fold change in band intensity compared to the CV. All raw audioradiograms are presented in **Tables S7-12**.

### ddPCR

Primers for PSI (HEX, Bio-Rad custom Assay, Hercules, California) were used to detect viral integration and PCBP2 (FAM, Bio-Rad custom Assay) as an integrated control in mouse cells and RPP30 (FAM, Bio-Rad custom Assay, Hercules, California) in the human fibroblast cells. Reactions were done with ddPCR Supermix for probes (no dUTP) (Bio-Rad, Hercules, California). Primers are detailed in **Table S5**. ddPCR plates were prepared using a Bio-Rad automated droplet generator and Px1 plate sealer. The PCR was conducted on a Bio-Rad C1000 thermocycler and analyzed on the Bio-Rad QX200.

### Acetone precipitation and conditioned media transfer

We conducted a TCA-DOC culture media protein precipitation according to Pare et al^40^ to assess for media secretion of ARSA. After 72 hours of culture, acetone precipitation was conducted on media harvested from these cells to precipitate all secreted proteins. The concentrated media preparation was run on a western blot and probed for ARSA.

Conditioned media was generated by incubating EA1 transduced fibroblasts in growth media for 72 hours. The conditioned media was centrifuged at 300xg for 5 minutes to precipitate any left-over cells or cellular debris. The supernatant was harvested for ARSA activity assay or incubated with untransduced fibroblasts for 48 hours. Harvested conditioned media was concentrated using a Centricon® plus centrifugal filter for 100Kda before measuring activity. Conditioned media-incubated fibroblasts were then PBS washed 2x after media incubation, then cell pellets were harvested and lysed for ARSA activity levels as described by the ARSA activity assay.

### EV enrichment and purification

EVs from transduced fibroblast cultures were harvested using the previously described ExtraPEG method.^41^ Briefly, cell-conditioned media was centrifuged at 500 × g for 5 min, 2,000 × g for 10 min, and then 10,000 (10K) × g for 30 min before incubating with a 1:1 volume of 2× PEG solution (16%, wt/vol, polyethylene glycol, 1 M NaCl) overnight. Solutions were centrifuged for one hour at 100,000 (100K) × g to obtain crude EV pellets. Pellets from 10K x g and 100K x g spins were lysed in 2× nonreducing Laemmli sample buffer (4% SDS, 100 mM Tris-HCl [pH 6.8], 0.4 mg/ml bromophenol blue, 20% glycerol) for immunoblot analysis. For further purification of small EVs, pellets from 100K x g spins were resuspended in 1.5 mL of 0.25 M sucrose buffer [10 mM Tris, pH 7.4] for bottom-loaded floatation gradient separation. Briefly, 60% iodixanol (Optiprep; Sigma, D1556, St. Louis, MO) was added 1:1 to the EV/buffer solution and transferred to the bottom of a 5.5 mL ultracentrifugation tube. Subsequently, 1.3 mL of 20% iodixanol and 1.2 mL of 10% iodixanol were carefully layered on top. Gradients were ultracentrifuged at maximum speed in an SW-55 rotor for 70 minutes with minimum deceleration break. Ten fractions of 490 μL were taken from the top of the gradient. Fractions 3, 5, and 6, previously demonstrated to contain purified EVs^42,28^, were diluted in PBS and washed by ultracentrifugation at 100,000 x g for 2 hours before lysis in 2× nonreducing Laemmli sample buffer for immunoblot analysis. We have previously published extensive and reproducible characterization of purified vesicles following these detailed approaches according to minimal information for studies of extracellular vesicles (MISEV) 2018 guidelines.^41,29,43,28^

### Long-term replating assay

Transduction of HSCs was performed to achieve VCNs in the range of 4-6 to maximize transformation. Following 2-3 weeks of expansion post-transduction to achieve millions of cells for replating, cells were plated into multi-well culture dishes and replated as the proliferation of the cells dictated. Transformed cells grow and consume media, while cells in which genomic mutagenesis did not occur enter senescence and stop growing. Mouse HSC cells were transduced with a pMSCV-GFP vector as a positive control of cell transformation. In contrast, cells transduced with EA1, EA2, and CV demonstrated no immortalization of mouse HSCs after nine weeks in culture. This was indicated by a lack of media depletion (and confirmed by microscopy) with VCN ranging from 1.0 to 6.3.

### Colony Forming Unit (CFU) Assay

Whole bone marrow cells were isolated from WT-transduced and WT-untransduced harvested mouse bones. Whole bone marrow cells were suspended in MethoCult GF M3534 (StemCell Technologies, Vancouver, Canada) as indicated by the manufacturer. Briefly, cells were resuspended in a 3mL thawed aliquot of MethoCult GF M3534. After preparation in the 3mL syringe with the 16-gauge blunt-end needles (28110), 1.1 mL of Cells were dispensed per well into a 6-well SmartDish to a final concentration of 50,000 cells per assay well. The myeloid population was interesting for CFU assay because it is the progenitor population crossing the BBB after BMT to generate microglia. Each condition was plated in duplicate according to the MethoCult technical guide. The assay measures the number of colonies formed in the myeloid-defined media after two weeks in culture using the stem vision plate analyzer (Stem Cell Vision, Stem Cell Technologies, Vancouver, Canada).

### Mice, Lineage Negative selection and mouse bone marrow transplantation

*Arsa*-KO (Cyagen, Santa Clara, CA). GFP or wild-type (WT) were purchased from the Jackson Laboratory (Bar Harbor, ME, USA) CD45.2 mice were bred at the Children’s Hospital of Philadelphia (CHOP) facility. Lin^-^ cells isolated from the bone marrow of donor mice (8–12 weeks old) via immunomagnetic separation using a mouse lineage cell depletion kit (Miltenyi Biotec, Auburn, CA, USA). Lin^-^ cells were transduced overnight at a viral multiplicity of infection (MOI) of 0.3-50 (**Table S6)** in StemSpan Serum Free Expansion Medium (SFEM) culture medium (StemCell Technologies, Vancouver, Canada) supplemented with 50 ng/mL recombinant murine stem cell factor (SCF), 10 ng/mL recombinant murine interleukin-6 (IL-6), 6 ng/mL IL-3 (PeproTech, Rocky Hill, NJ, USA), 200 mM L-glutamine, 100 U/mL penicillin/ streptomycin (Gibco, Thermo Fisher Scientific, Waltham, MA), and 2 uL Lentiblast Premium (OZBiosciences, San Diego, CA). Specific cell numbers and MOI for CV, EA1 and EA2 Arsa-KO mouse transplants are found in **Table S5**. A minimum of 300,000 viable Lin^-^ donor cells in phosphate-buffered saline (PBS) (Gibco, Thermo Fisher Scientific, Waltham, MA). were intravenously injected into lethally irradiated ARSA KO recipient mice (8–12 weeks old) of the opposite gender. For WT mice, 2 rounds of virus transduction, 18 hours apart at MOI between ∼10-50.

### Behavior assays

Rotarod assays were started with a training period two days before the assay time. Mice were trained at 10 RPM for 300 seconds. Mice were placed back on the rod if a fall occurred to acclimatize them. Three trials were done with a 15-minute break between each trial. The trial began two days after the training period and was conducted with a 5-40 RPM ramp over 300 seconds. This was done for three trials with a 15-minute break between 5 days in a row. Pole descent assay was conducted using a ring stand rod taped and covered with a cloth lab sheet for grip. The top was covered with a large spherical object to prevent mice from going to the top and hugging the pole. The tape was placed as a starting region for the first 5-6 cm. The distance from the tape start to the bottom was approximately 34 cm. An acclimation period was conducted for the mice to familiarize them with the space and rod until a basic proficiency for descent was reached. All animals were video recorded during descent; times and loss of footing/traction were assessed from the recording after passing the marked tape start line. The pole descent assay metrics were split into two parameters: time of descent over a specified pole distance and loss of traction events for the mice during descent. A traction loss event was defined as a sudden and uncontrolled fall, drop, or slip downwards during pole descent, reported as events per trail. Descent time was evaluated by measuring the uninterrupted downward movement of the mice across half of a section of the 34 cm traversed (approximately 17cm). Change of direction, upward movement, or stopping on the pole was not part of the descent duration measure. This ensured the highest fidelity and robustness of measured descent data, accumulating more measures of descent time per trial and eliminating apparent/uncontrolled movement and variability from assessed times.

### Flow Cytometry

Flow cytometry was performed using a BD FACSCalibur (BD Biosciences, Franklin Lakes, NJ) instrument. Erythroid populations were characterized using FITC Anti-Mouse CD71 (113806) and APC Anti-Mouse Ter119 (116212), PE Anti-Human/Mouse CD44 (103008), as well as 7-AAD Viability Staining Solution (420404) (Biolegend, San Diego, CA). Lymphoid populations were characterized using APC anti-mouse CD3 (100236, Biolegend, San Diego, CA) and PE-Cyanine7 CD45R (B220) Monoclonal Antibody (RA3-6B2) (25045282, ThermoFisher Scientific, Waltham, MA). The flow cytometry results were analyzed using FlowJo™ v10.8 Software (BD Life Sciences).

### Histological Processing

Mice were anesthetized and transcardially perfused with an initial flush of PBS followed by 4% paraformaldehyde (PFA). Anesthetic was provided by weight-based injection of a mixture of ketamine/xylazine for adult mice (>P21) with a supplemental isoflurane/oxygen flow. The harvested whole brain tissue was post-fixed with 4% PFA in PBS overnight at 4°C, then rinsed in 1X Phosphate buffer saline (PBS; Thermo Scientific) and transferred to 30% sucrose in PBS at 4°C until the sample was no longer buoyant (>3 days), indicating penetrance of the sucrose solution. The whole brain was then embedded in a Tissue-Tek Optimum Cutting Temperature compound (Sakura, 4583) and cryosectioned by a cryostat microtome (Leica, CM3050S) into 50µm sections in a coronal orientation. Sections were stored in free-floating in PBS and transferred to a freezing medium comprised of a mixture of DMEM containing HEPES (Gibco, 11885084 and Sigma, H7523, respectively) and glycerol (Sigma, G6279-500) for long-term storage at -20°C.

### Staining

#### Alcian Blue

Alcian blue solution was prepared using 3% glacial acetic acid (Amresco, 0714-500) and Alcian blue 8GX (Sigma, A-5268) with a pH adjusted to 2.5 and filtration after overnight stirring. Brain sections of 50µm thickness were mounted onto a Superfrost Plus slide glass (Fisher, 12-550-15) and dried at room temperature. These slides were then quickly hydrated in distilled water, submerged in 3% acetic acid, and incubated in Alcian blue solution for 1 hour at room temperature. Sections were washed in tap water and rinsed in distilled water before exposure to Nuclear Fast Red Counterstain (Vector,

H-3403). After washing in tap water, slides were dehydrated in increasing ethanol gradients, cleared with Histo-Clear (National Diagnostics, HS-200), and cover-slipped with ThermoFisher Gold Seal Cover Glass 24 x 60mm No.1 thickness (Electron Microscopy Sciences, 63770-01). Sections were imaged using a Leica CTR6000/B microscope to quantify sulfatide accumulation in different brain areas.

#### Eriochrome Cyanine

Eriochrome Cyanine (Eri-C) staining was conducted using previously established methods.^44^ Sections were imaged using a Leica CTR6000/B microscope, and Image J software was used to quantify demyelination vacuoles in the corpus callosum.

#### Immunofluorescence

Brain sections of 50µm thickness were washed in PBS, blocked in 10% Normal Goat Serum (EMD Millipore, S26), 2% Bovine Serum Albumin (Sigma, A7030-100G), and 0.1% Triton X-100 (Sigma, T-9284) in PBS for 1 hour at room temperature, and incubated in primary antibodies (See **Antibody Resource Table**) overnight at 4°C. Following this incubation, sections were washed with 0.1% Triton X-100 (Sigma, T-9284) in PBS, incubated in fluorescent secondary antibodies for 1 hour, and protected from light at room temperature. The secondary antibodies used in this study are AlexaFluor-488, AlexaFluor-555, and AlexaFluor-647 conjugated secondary antibodies against rat (Invitrogen, A21208), mouse (Invitrogen, A315570), and rabbit (Invitrogen, A21245) at 1:1000 dilutions. Sections were sequentially washed with 0.1% Triton X-100 (Sigma, T-9284) in PBS, counterstained with DAPI, and mounted onto Superfrost Plus Microscope Slides (Fisher, 12-550-15). The tissues were mounted using ProLong Gold Antifade Mountant with DAPI (ThermoFisher, P36931). Sections were imaged using a Keyence BZ-X810 microscope and a Keyence BZ-X800 Analyzer system to quantify the immunofluorescence staining.

### Antibody Resource Table

| Antibody | Type/<br>Species | Source | Identifiers | Additional<br>Information |
| --- | --- | --- | --- | --- |
| Anti-GFAP | Rat | IDDRC hybridoma,<br>courtesy Dr. Judith<br>Grinspan | - | IHC Dilution 1:1 |
| Anti-Iba1 | Rabbit<br>polyclonal | FUJIFILM Wako Pure<br>Chemical Corporation | 019-19741 | IHC Dilution 1:250 |
| Anti-NeuN | Mouse<br>monoclonal | Millipore Sigma | MAB377 | IHC Dilution 1:250 |

### Statistics

All statistical analysis was performed using Prism 10 for MacOS (GraphPad Software, Boston, Massachusetts USA, www.graphpad.com, Software, San Diego, USA). A comparison of two independent groups was conducted by unpaired t-test. A one-way ANOVA was used to compare more than two groups. Rotarod data were presented as SEM. The rest of the data were represented as mean ± standard deviation. Significance levels were *P<0.05, **P<0.01, ***P<0.001 and ****P<0.0001. If no statistical significance was found, it was not reported.

### Institutional Animal Care and Use Committee (IACUC) regulation

All bone marrow transplantation studies in mice were approved by the IACUC (protocol #1173) at the CHOP. Reporting of animal studies has been provided per ARRIVE guidelines.

### Data Sharing Statement

Additional data supporting this study’s findings are available on request from the corresponding author. No public repository data have been generated for this publication.

### COI

S.R. is a scientific advisory board member of Ionis Pharmaceuticals, Vifor, and Disc Medicine. Present-last five years: S.R. has been or is a consultant for GSK, BMS, Incyte, Cambridge Healthcare Res, Celgene Corporation, Catenion, First Manhattan Co., FORMA Therapeutics, Ghost Tree Capital, Keros Therapeutics, Noble insight, Protagonist Therapeutics, Sanofi Aventis U.S., Slingshot Insight, Spexis AG, Techspert.io, BVF Partners L.P., Rallybio, LLC, venBio Select LLC, ExpertConnect LLC, LifeSci Capital.

### Support

Digestive and Kidney Diseases Institute of the National Institutes of Health (R01 DK133475, R01 DK095112), Institute for Translational Medicine and Therapeutics (ITMAT), Irish Health Research Board-Health Research Charities Ireland (HRCI-HRB), Acceleration-Seed program/CHOP and The Sickle Cell and Red Cell Disorders Curative Therapy Center (CuRED)-Frontier Program & Molecular Therapies for Inborn Errors of Metabolism-Frontier Program to S.R. R01-HL164633 grant to P.K. and grant R38-HL143613-04 to S.N.H. Calliope Joy Foundation, to S.R and A.V. Kamens chair in translational neurotherapeutics to A.V.

## ACKNOWLEDGMENTS

David Wenger from Thomas Jefferson University was instrumental in training us on the ARSA colorimetric enzymatic assay, allowing us to adapt it for our laboratory use in 96 well plate format.

## REFERENCES

1. Rosenberg, J.B., Kaminsky, S.M., Aubourg, P., Crystal, R.G., and Sondhi, D. (2016). Gene therapy for metachromatic leukodystrophy. J Neurosci Res 94, 1169–1179. 10.1002/jnr.23792.

2. Brimley, C.J., Lopez, J., van Haren, K., Wilkes, J., Sheng, X., Nelson, C., Korgenski, E.K., Srivastava, R., and Bonkowsky, J.L. (2013). National variation in costs and mortality for leukodystrophy patients in US children’s hospitals. Pediatr Neurol 49, 156–162 e151. 10.1016/j.pediatrneurol.2013.06.006.

3. Papaioannou, I., Owen, J.S., and Yanez-Munoz, R.J. (2023). Clinical applications of gene therapy for rare diseases: A review. Int J Exp Pathol 104, 154–176. 10.1111/iep.12478.

4. Matzner, U., Breiden, B., Schwarzmann, G., Yaghootfam, A., Fluharty, A.L., Hasilik, A., Sandhoff, K., and Gieselmann, V. (2009). Saposin B-dependent reconstitution of arylsulfatase A activity in vitro and in cell culture models of metachromatic leukodystrophy. J Biol Chem 284, 9372–9381. 10.1074/jbc.M809457200.

5. Schmidt, J.L., Pizzino, A., Nicholl, J., Foley, A., Wang, Y., Rosenfeld, J.A., Mighion, L., Bean, L., da Silva, C., Cho, M.T., Truty, R., et al. (2020). Estimating the relative frequency of leukodystrophies and recommendations for carrier screening in the era of next-generation sequencing. Am J Med Genet A 182, 1906–1912. 10.1002/ajmg.a.61641.

6. Farah, M.H., Dali, C.I., Groeschel, S., Moldovan, M., Whiteman, D.A.H., Malanga, C.J., Krageloh-Mann, I., Li, J., Barton, N., and Krarup, C. (2024). Effects of sulfatide on peripheral nerves in metachromatic leukodystrophy. Ann Clin Transl Neurol 11, 328–341. 10.1002/acn3.51954.

7. Fumagalli, F., Calbi, V., Natali Sora, M.G., Sessa, M., Baldoli, C., Rancoita, P.M.V., Ciotti, F., Sarzana, M., Fraschini, M., Zambon, A.A., Acquati, S., et al. (2022). Lentiviral haematopoietic stem-cell gene therapy for early-onset metachromatic leukodystrophy: long-term results from a non-randomised, open-label, phase 1/2 trial and expanded access. Lancet 399, 372–383. 10.1016/S0140-6736(21)02017-1.

8. Biffi, A., Capotondo, A., Fasano, S., del Carro, U., Marchesini, S., Azuma, H., Malaguti, M.C., Amadio, S., Brambilla, R., Grompe, M., Bordignon, C., et al. (2006). Gene therapy of metachromatic leukodystrophy reverses neurological damage and deficits in mice. J Clin Invest 116, 3070–3082. 10.1172/JCI28873.

9. Biffi, A., Montini, E., Lorioli, L., Cesani, M., Fumagalli, F., Plati, T., Baldoli, C., Martino, S., Calabria, A., Canale, S., Benedicenti, F., et al. (2013). Lentiviral hematopoietic stem cell gene therapy benefits metachromatic leukodystrophy. Science 341, 1233158. 10.1126/science.1233158.

10. Sessa, M., Lorioli, L., Fumagalli, F., Acquati, S., Redaelli, D., Baldoli, C., Canale, S., Lopez, I.D., Morena, F., Calabria, A., Fiori, R., et al. (2016). Lentiviral haemopoietic stem-cell gene therapy in early-onset metachromatic leukodystrophy: an ad-hoc analysis of a non-randomised, open-label, phase 1/2 trial. Lancet 388, 476–487. 10.1016/S0140-6736(16)30374-9.

11. Biffi, A., De Palma, M., Quattrini, A., Del Carro, U., Amadio, S., Visigalli, I., Sessa, M., Fasano, S., Brambilla, R., Marchesini, S., Bordignon, C., et al. (2004). Correction of metachromatic leukodystrophy in the mouse model by transplantation of genetically modified hematopoietic stem cells. J Clin Invest 113, 1118–1129. 10.1172/JCI19205.

12. Ginhoux, F., and Prinz, M. (2015). Origin of microglia: current concepts and past controversies. Cold Spring Harb Perspect Biol 7, a020537. 10.1101/cshperspect.a020537.

13. Schoenmakers, D.H., Mochel, F., Adang, L.A., Boelens, J.J., Calbi, V., Eklund, E.A., Gronborg, S.W., Fumagalli, F., Groeschel, S., Lindemans, C., Sevin, C., et al. (2024). Inventory of current practices regarding hematopoietic stem cell transplantation in metachromatic leukodystrophy in Europe and neighboring countries. Orphanet J Rare Dis 19, 46. 10.1186/s13023-024-03075-3.

14. Kolata, G. (2021). Sickle cell gene therapy trial halted after two patients develop cancer, though the link is not certain. Genetic Literacy Project.

15. MARAC (2021). Temporary Suspension of Clinical Trials.

16. Breda, L., Ghiaccio, V., Tanaka, N., Jarocha, D., Ikawa, Y., Abdulmalik, O., Dong, A., Casu, C., Raabe, T.D., Shan, X., Danet-Desnoyers, G.A., et al. (2021). Lentiviral vector ALS20 yields high hemoglobin levels with low genomic integrations for treatment of beta-globinopathies. Mol Ther 29, 1625–1638. 10.1016/j.ymthe.2020.12.036.

17. Zufferey, R., Donello, J.E., Trono, D., and Hope, T.J. (1999). Woodchuck hepatitis virus posttranscriptional regulatory element enhances expression of transgenes delivered by retroviral vectors. J Virol 73, 2886–2892. 10.1128/JVI.73.4.2886-2892.1999.

18. Rivella, S., Callegari, J.A., May, C., Tan, C.W., and Sadelain, M. (2000). The cHS4 insulator increases the probability of retroviral expression at random chromosomal integration sites. J Virol 74, 4679–4687. 10.1128/jvi.74.10.4679-4687.2000.

19. Browning, D.L., Everson, E.M., Leap, D.J., Hocum, J.D., Wang, H., Stamatoyannopoulos, G., and Trobridge, G.D. (2017). Evidence for the in vivo safety of insulated foamy viral vectors. Gene Ther 24, 187–198. 10.1038/gt.2016.88.

20. Browning, D.L., and Trobridge, G.D. (2016). Insulators to Improve the Safety of Retroviral Vectors for HIV Gene Therapy. Biomedicines 4. 10.3390/biomedicines4010004.

21. Romero, Z., Campo-Fernandez, B., Wherley, J., Kaufman, M.L., Urbinati, F., Cooper, A.R., Hoban, M.D., Baldwin, K.M., Lumaquin, D., Wang, X., Senadheera, S., et al. (2015). The human ankyrin 1 promoter insulator sustains gene expression in a beta-globin lentiviral vector in hematopoietic stem cells. Mol Ther Methods Clin Dev 2, 15012. 10.1038/mtm.2015.12.

22. Browning, D.L., Collins, C.P., Hocum, J.D., Leap, D.J., Rae, D.T., and Trobridge, G.D. (2016). Insulated Foamy Viral Vectors. Hum Gene Ther 27, 255–266. 10.1089/hum.2015.110.

23. Kingsman, S.M., Mitrophanous, K., and Olsen, J.C. (2005). Potential oncogene activity of the woodchuck hepatitis post-transcriptional regulatory element (WPRE). Gene Ther 12, 3–4. 10.1038/sj.gt.3302417.

24. Skarpidi, E., Vassilopoulos, G., Stamatoyannopoulos, G., and Li, Q. (1998). Comparison of expression of human globin genes transferred into mouse erythroleukemia cells and in transgenic mice. Blood 92, 3416–3421.

25. Domingues, H.S., Portugal, C.C., Socodato, R., and Relvas, J.B. (2016). Oligodendrocyte, Astrocyte, and Microglia Crosstalk in Myelin Development, Damage, and Repair. Front Cell Dev Biol 4, 71. 10.3389/fcell.2016.00071.

26. Wolf, N.I., Breur, M., Plug, B., Beerepoot, S., Westerveld, A.S.R., van Rappard, D.F., de Vries, S.I., Kole, M.H.P., Vanderver, A., van der Knaap, M.S., Lindemans, C.A., et al. (2020). Metachromatic leukodystrophy and transplantation: remyelination, no cross-correction. Ann Clin Transl Neurol 7, 169–180. 10.1002/acn3.50975.

27. Do, M.A., Levy, D., Brown, A., Marriott, G., and Lu, B. (2019). Targeted delivery of lysosomal enzymes to the endocytic compartment in human cells using engineered extracellular vesicles. Sci Rep 9, 17274. 10.1038/s41598-019-53844-5.

28. Hurwitz, S.N., Olcese, J.M., and Meckes, D.G., Jr. (2019). Extraction of Extracellular Vesicles from Whole Tissue. J Vis Exp. 10.3791/59143.

29. Hurwitz, S.N., and Meckes, D.G., Jr. (2017). An Adaptable Polyethylene Glycol-Based Workflow for Proteomic Analysis of Extracellular Vesicles. Methods Mol Biol 1660, 303–317. 10.1007/978-1-4939-7253-1_25.

30. Ikawa, Y., Uchiyama, T., Jagadeesh, G.J., and Candotti, F. (2016). The long terminal repeat negative control region is a critical element for insertional oncogenesis after gene transfer into hematopoietic progenitors with Moloney murine leukemia viral vectors. Gene Ther 23, 815–818. 10.1038/gt.2016.51.

31. Ory, D.S., Neugeboren, B.A., and Mulligan, R.C. (1996). A stable human-derived packaging cell line for production of high titer retrovirus/vesicular stomatitis virus G pseudotypes. Proc Natl Acad Sci U S A 93, 11400–11406. 10.1073/pnas.93.21.11400.

32. Matzner, U., and Gieselmann, V. (2005). Gene therapy of metachromatic leukodystrophy. Expert Opin Biol Ther 5, 55–65. 10.1517/14712598.5.1.55.

33. Servick, K. (2021). Gene therapy clinical trial halted as cancer risk surfaces. Science.

34. Ferreira, M.V., Cabral, E.T., and Coroadinha, A.S. (2021). Progress and Perspectives in the Development of Lentiviral Vector Producer Cells. Biotechnol J 16, e2000017. 10.1002/biot.202000017.

35. Thielen, F.W., Heine, R., Berg, S.V.D., Ham, R., and Groot, C.A.U. (2022). Towards sustainability and affordability of expensive cell and gene therapies? Applying a cost-based pricing model to estimate prices for Libmeldy and Zolgensma. Cytotherapy 24, 1245–1258. 10.1016/j.jcyt.2022.09.002.

36. Cornetta, K., Bonamino, M., Mahlangu, J., Mingozzi, F., Rangarajan, S., and Rao, J. (2022). Gene therapy access: Global challenges, opportunities, and views from Brazil, South Africa, and India. Mol Ther 30, 2122–2129. 10.1016/j.ymthe.2022.04.002.

37. Jelin, A.C., and Vora, N. (2018). Whole Exome Sequencing: Applications in Prenatal Genetics. Obstet Gynecol Clin North Am 45, 69–81. 10.1016/j.ogc.2017.10.003.

38. Dukhovny, S., and Norton, M.E. (2018). What are the goals of prenatal genetic testing? Semin Perinatol 42, 270–274. 10.1053/j.semperi.2018.07.002.

39. Baum, H., Dodgson, K.S., and Spencer, B. (1959). The assay of arylsulphatases A and B in human urine. Clin Chim Acta 4, 453–455. 10.1016/0009-8981(59)90119-6.

40. Pare, B., Deschenes, L.T., Pouliot, R., Dupre, N., and Gros-Louis, F. (2016). An Optimized Approach to Recover Secreted Proteins from Fibroblast Conditioned-Media for Secretomic Analysis. Front Cell Neurosci 10, 70. 10.3389/fncel.2016.00070.

41. Rider, M.A., Hurwitz, S.N., and Meckes, D.G., Jr. (2016). ExtraPEG: A Polyethylene Glycol-Based Method for Enrichment of Extracellular Vesicles. Sci Rep 6, 23978. 10.1038/srep23978.

42. Kowal, J., Arras, G., Colombo, M., Jouve, M., Morath, J.P., Primdal-Bengtson, B., Dingli, F., Loew, D., Tkach, M., and Thery, C. (2016). Proteomic comparison defines novel markers to characterize heterogeneous populations of extracellular vesicle subtypes. Proc Natl Acad Sci U S A 113, E968–977. 10.1073/pnas.1521230113.

43. Hurwitz, S.N., Sun, L., Cole, K.Y., Ford, C.R., 3rd, Olcese, J.M., and Meckes, D.G., Jr. (2018). An optimized method for enrichment of whole brain-derived extracellular vesicles reveals insight into neurodegenerative processes in a mouse model of Alzheimer’s disease. J Neurosci Methods 307, 210–220. 10.1016/j.jneumeth.2018.05.022.

44. Sase, S., Almad, A.A., Boecker, C.A., Guedes-Dias, P., Li, J.J., Takanohashi, A., Patel, A., McCaffrey, T., Patel, H., Sirdeshpande, D., Curiel, J., et al. (2020). TUBB4A mutations result in both glial and neuronal degeneration in an H-ABC leukodystrophy mouse model. Elife 9. 10.7554/eLife.52986.

