## Supplemental Figures and Tables for "Effective Gene Therapy for Metachromatic Leukodystrophy Achieved with Minimal Lentiviral Genomic Integrations"

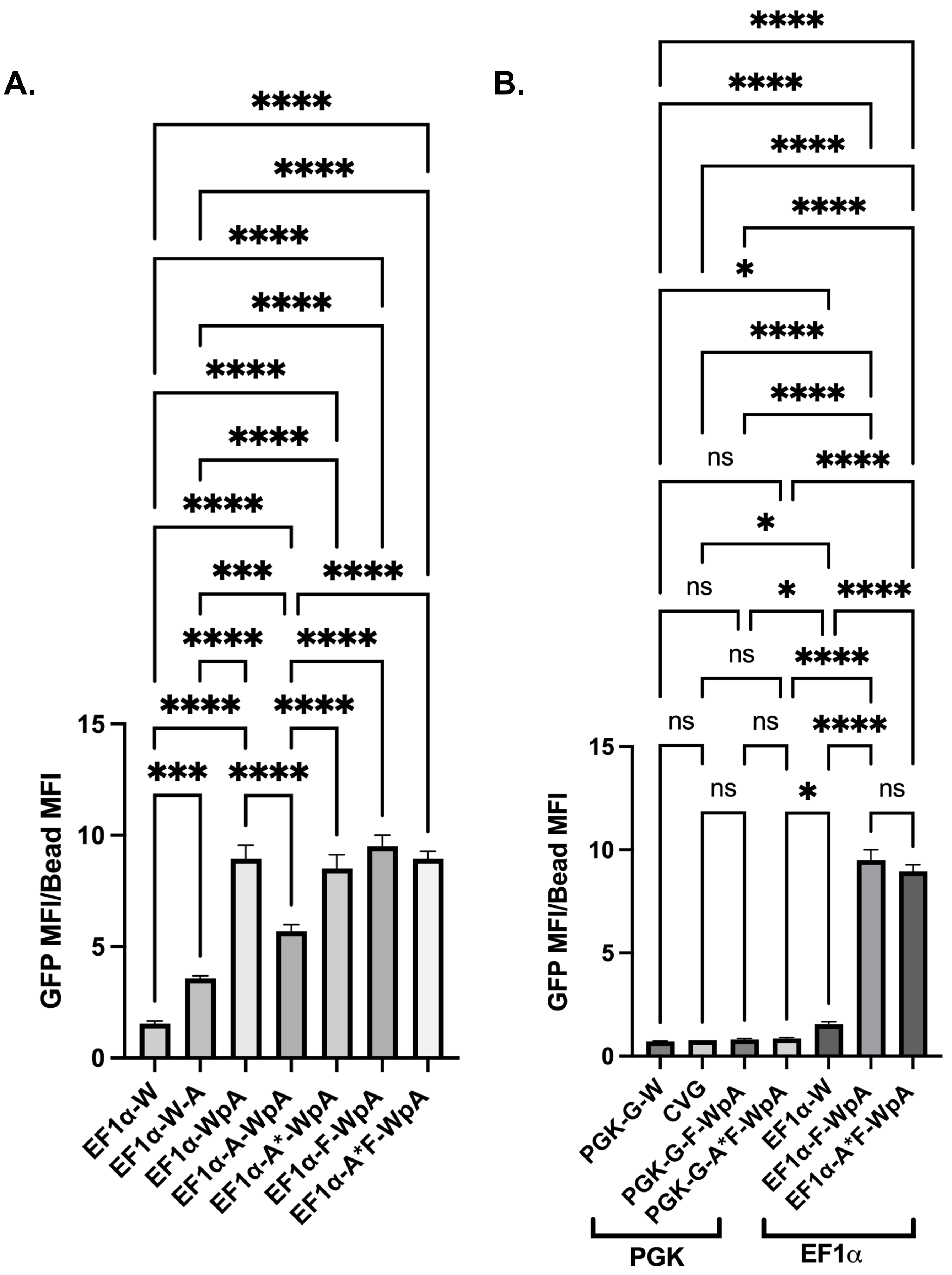

**S1. Complete representation of significant differences for Figure 1B and 1C.** (A) MFI for all EF1α vector arrangements shown above (n = 3). (B) MFI comparison of select PGK and the CVG arrangement compared to their EF1α counterparts. *P<0.05, **P<0.01, ***P<0.001, ****P<0.0001

### Supplemental Figure 1

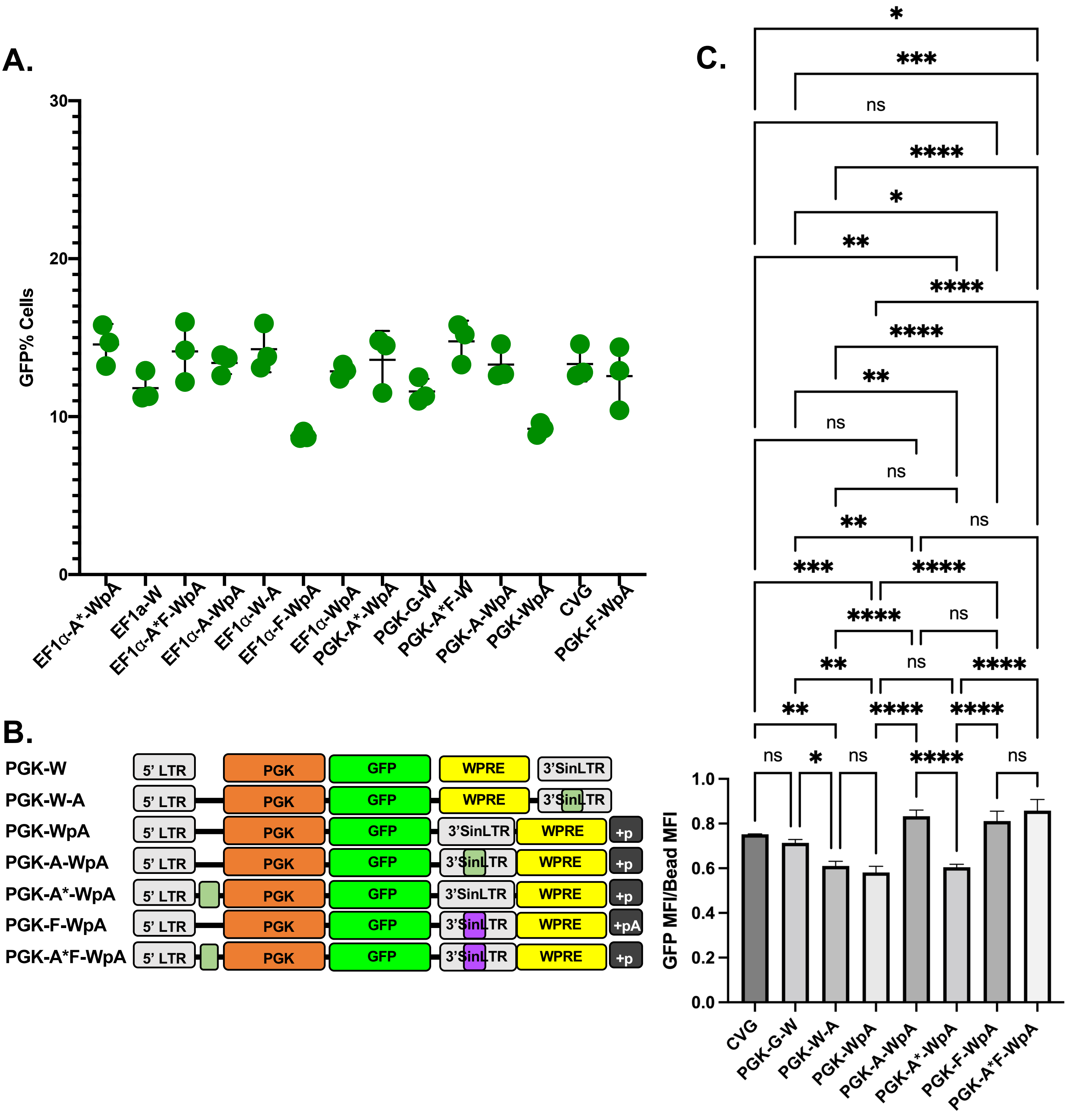

**Figure S2. GFP positive percentages of Mel cells transduced and complete representation of significant differences for all the PGK promoter-driven GFP reporter vectors. (**A) Associated GFP positivity percentages for each GFP test vector from **Figure 1A** in Mel cells. (B) Maps of the multiple vector arrangement used to determine the optimal configuration of PGK promoter with ankyrin or foamy insulators and WPRE (W) with PolyA tail (pA) region for optimal GFP MFI. (C) Complete representation of significant differences in all PGK driven promotors tested for Figure 1 (n = 3).

### Supplemental Figure 2

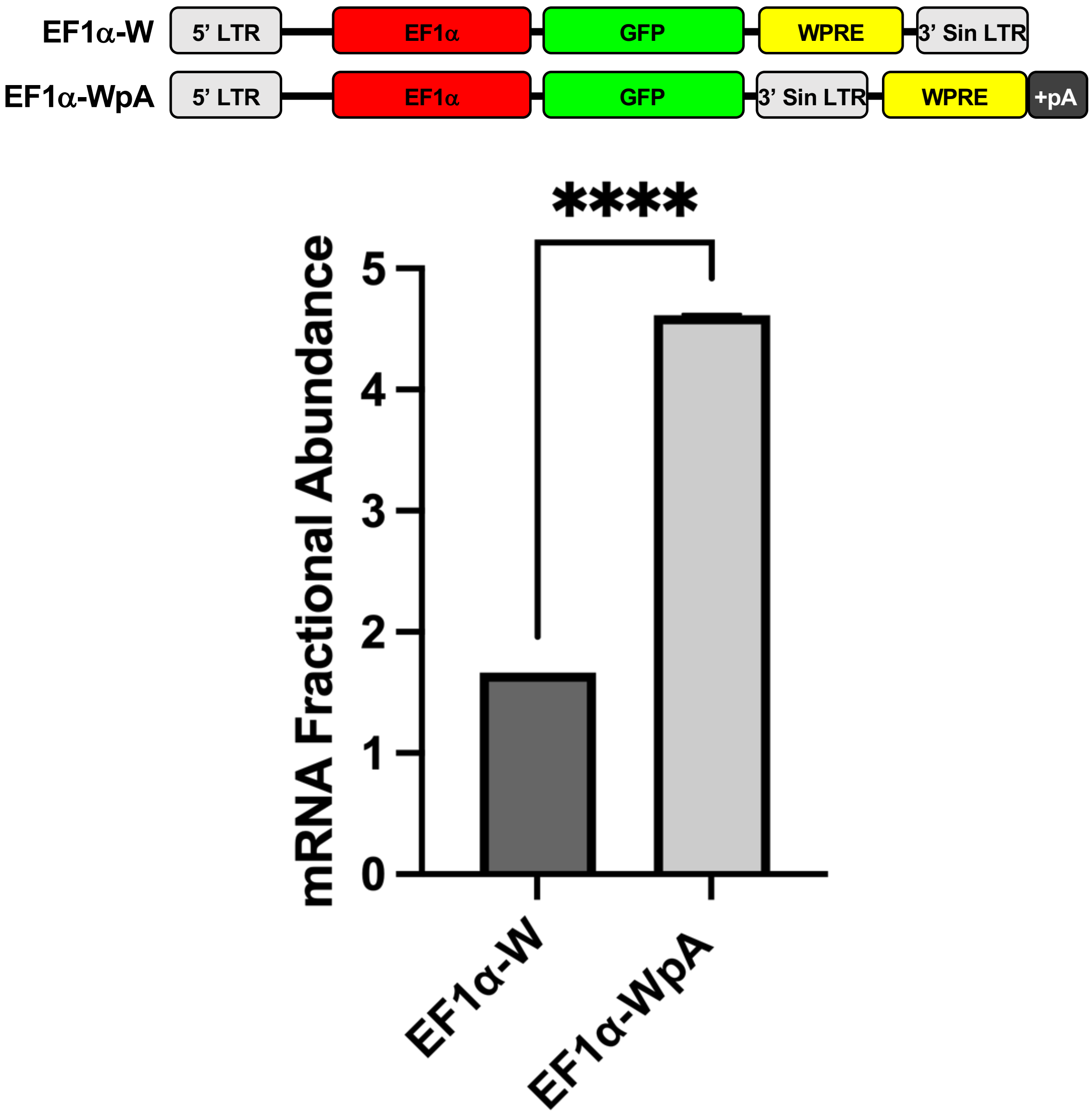

**S3. Expression of GFP mRNA in EF1α-driven vectors to assess the effect of WPRE rearrangement (n = 2).** mRNA expression of EF1α-W vs of EF1α –WpA by ddPCR.

#### Supplemental Figure 3

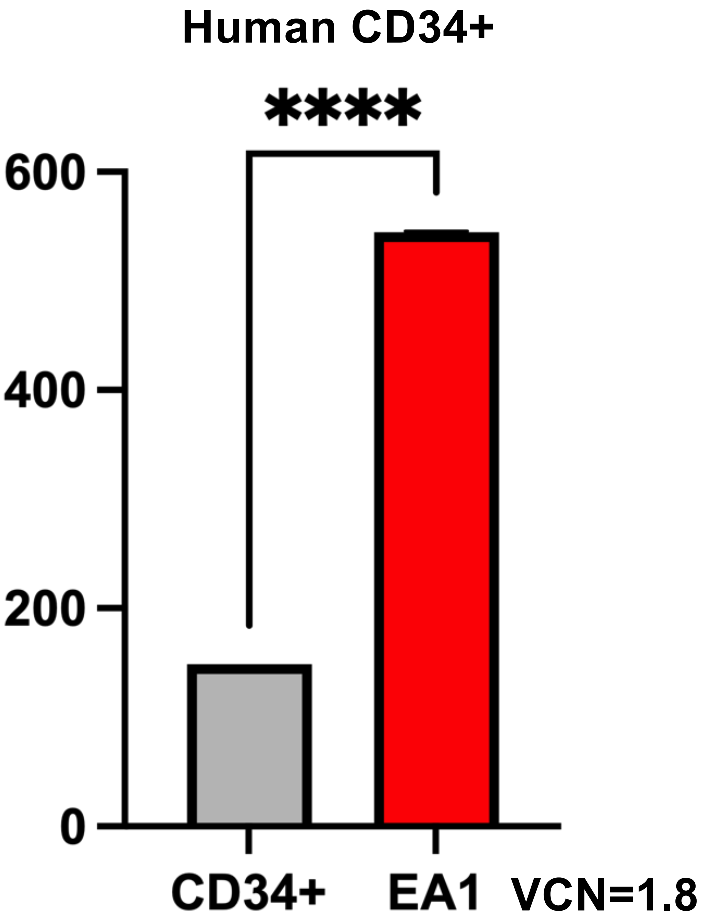

**S4. Vector EA1 transduced into human CD34+ bone marrow hematopoietic progenitor.** Cells demonstrating robust expression compared to naïve CD34+ cells (n = 2). This data indicates that the expression of EA1 is robust in CD34+ transduced cells, showing about 3x more ARSA expression than CD34+ cells alone. ARSA activity was normalized to VCN.

•Cell culture: CD34+ cells were thawed by rapidly incubating them at 37°C and then adding a 10X volume of IMDM 5% FBS drop by drop. Cells were first prestimulated in StemSpan SFEM (09650, STEMCELL Technologies, 1618 Station Street Vancouver, BC, V6A 1B6, Canada) supplemented with 300ng/mL hSCF (300-07, PEPRO TECH, 29 Margravine Road London W6 8LL, UK), 300ng/mL hFLT3 (300-19, PEPRO TECH) and 100ng/mL hTPO (300-18, PEPRO TECH). Cells were transduced with 10 MOI of EA1 2 times, 12 hours apart, using Poloxamer 338 2 µl/mL (P2164021, Sigma) and PGE2 0.141 µl/mL (72192, STEMCELL Technologies), as adjuvants. Cells were then subsequently cultured for 2 weeks in expansion media (StemSpan SFEM supplemented with CC-100 10ul/mL (02690 ,STEMCELL Technologies), Epogen (55513-126-10, Amgen, Thousand Oaks, California) 1uL/mL, Dexamethsone 1ul/mL (D2915, Sigma Aldrich) and Pen-Strep 10ul/mL(15140-122, LifeTechnologies, Carlsbad, CA) before harvest for ARSA activity assay.

#### Supplemental Figure 4

**A.**

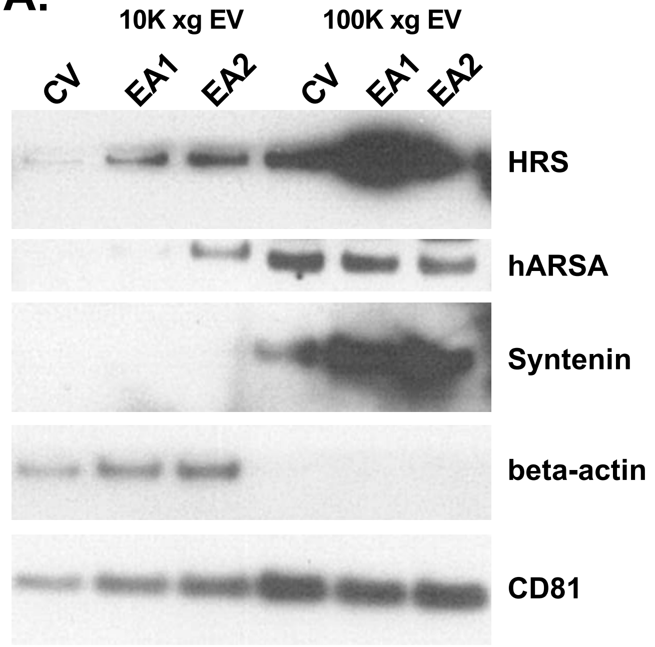

**B.**

| Sample | VCN |
|---|---|
| A. CV | 1.1 |
| B. EA1 | 0.54 |
| C. EA2 | 0.67 |

| Sample | Gradient Fractions |
|---|---|
| A. CV | A3, A5/6 |
| B. EA1 | B3 B5/6 |
| C. EA2 | C3 C5/6 |

**C.**

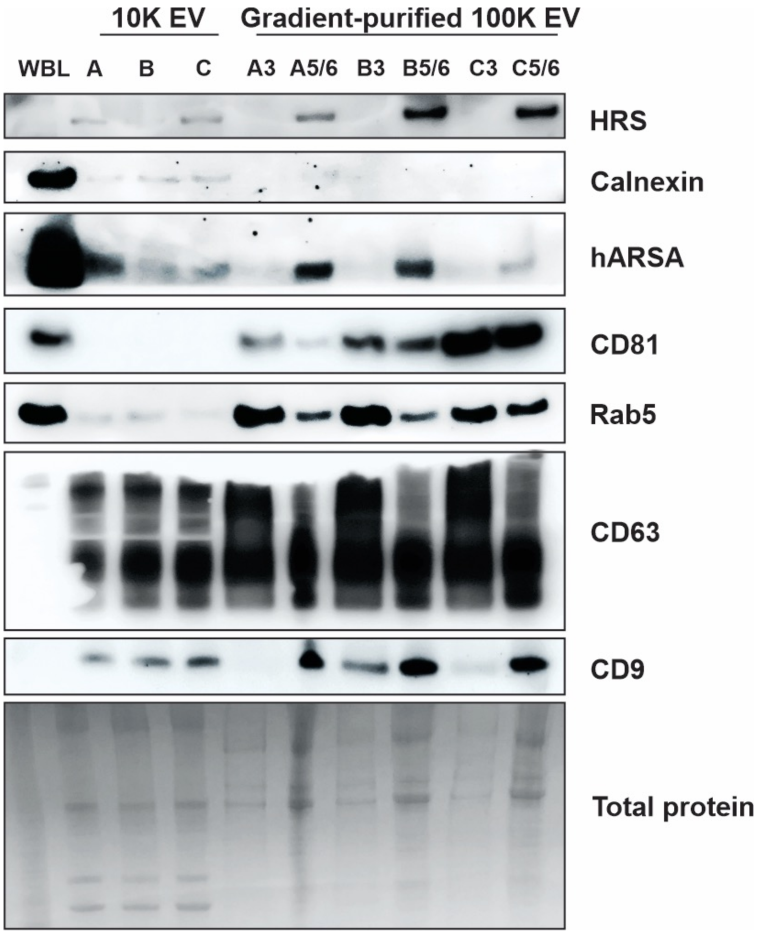

***WBL = WT mouse Brain/Cell Loading Control**

**Figure S5. ARSA protein secretion associated is associated with EVs. (**A) Western blot showing enrichment of ARSA in small EVs isolated from CV, EA1 and EA2-transduced cells in the 100K xg fraction compared to larger vesicles isolated in the 10K xg fraction. Enrichment of EV particles is evident by the enhanced presence of HRS, Syntenin and CD81, all markers for EVs. (B) Table indicating VCNs and gradient fractions in A for each treatment. (C) Western blot of 10K xg EV fraction and further iodixanol density gradient purified fractions 3, 5 and 6, which have previously demonstrated to be enriched in small EVs. The gradient purified fraction was depleted of ER protein Calnexin while enriched in numerous EV-associated proteins including CD9, CD63 and Rab5 (A GTPase associated with the plasma membrane and early endosomes).

#### Supplemental Figure 5

### Bone Marrow: Erythroid

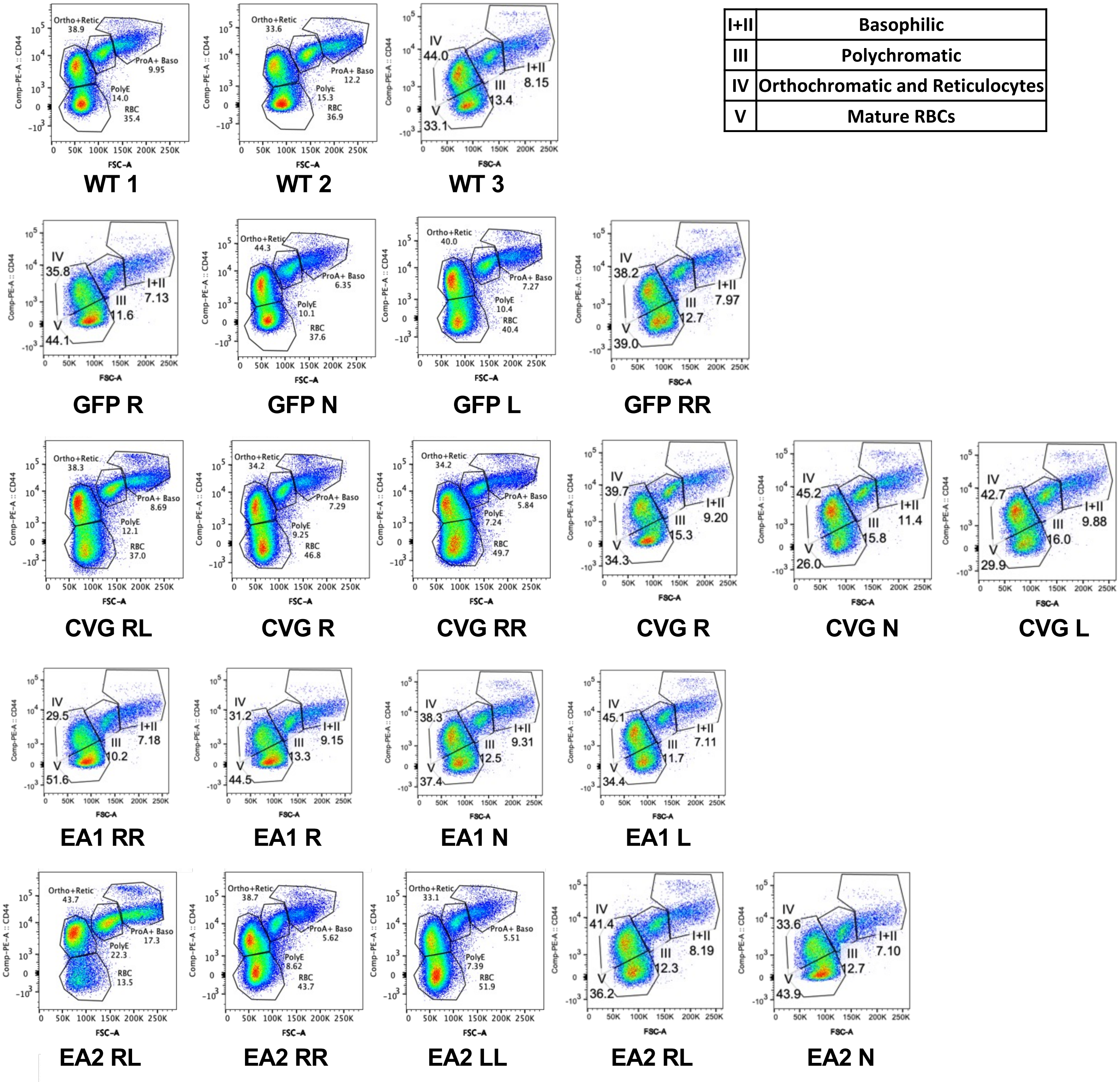

| | |
|---|---|
| I+II | Basophilic |
| III | Polychromatic |
| IV | Orthochromatic and Reticulocytes |
| V | Mature RBCs |

**Figure S6A. Extended analysis for vector transplanted WT mice from Figure 5.** Erythropoiesis analysis (population I to V, from immature to enucleated red cells) by flow cytometry in WT controls, GFP mice transplanted with normal marrow or mice treated with CVG, EA1 , or EA2, from top to bottom.

#### Supplemental Figure 6A

### Bone Marrow:
### Lymphocyte

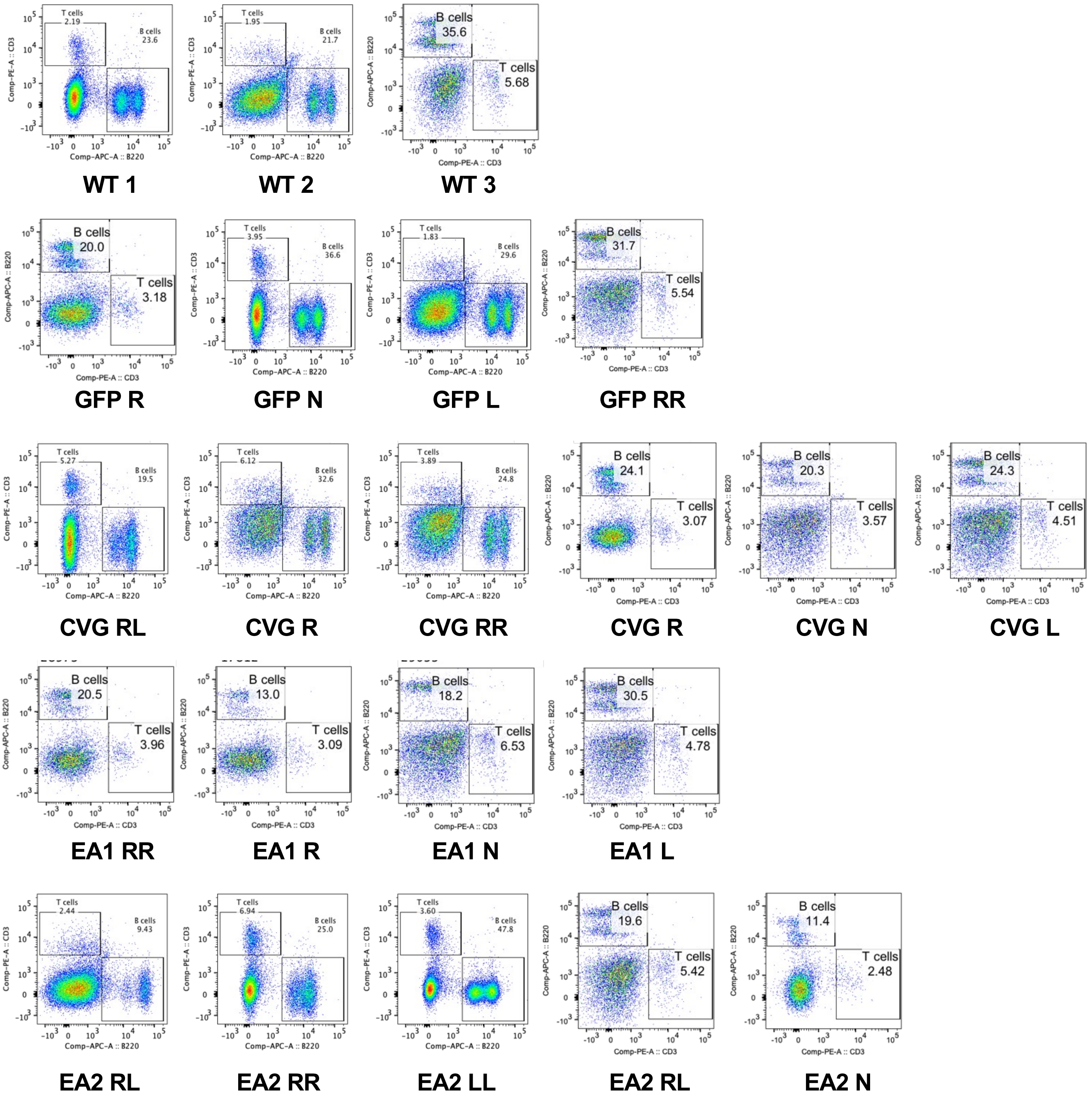

**Figure S6B. Extended analysis for vector transplanted WT mice from Figure 5.** Flow sort panel of lymphocyte B and T cells in the same animals as in S6A.

#### Supplemental Figure 6B

#### 1. Method

The gRNA to mouse Arsa gene, and Cas9 mRNA were co-injected into fertilized mouse eggs to generate targeted knockout offspring. F0 founder animals were identified by PCR followed by sequence analysis, which were bred to wildtype mice to test germline transmission and F1 animal generation. Inter-cross heterozygous F1 mice to generate homozygous F2 mice.

#### 2. gRNA target sequence

gRNA1 (matching reverse strand of gene): GGCAGTCTCGGATACGCCCGCGG
gRNA2 (matching reverse strand of gene): TGGTAAGGTGGCATCGGACTTGG

#### 3. Targeting Strategy

**The (start codon) ATG is 933 bp into exon 2 of the mouse *Arsa* gene**

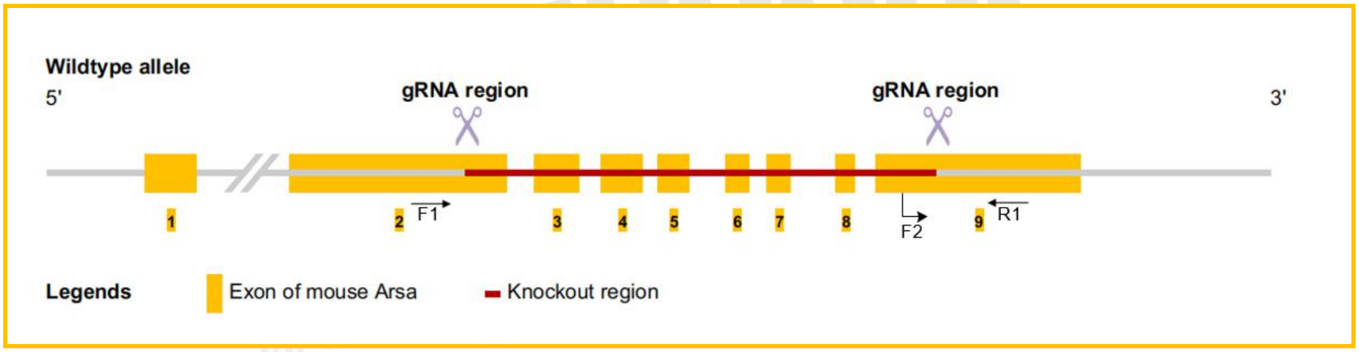

#### 4. Genotyping strategy

Primers1: (Annealing Temperature 60.0 ºC)

F1: 5'-AAGAACGTAAGGTTACCAAGCCC-3'
R1: 5'-AGGTCCTTTAACTCTACCCGGAA-3'
Product size: 610 bp

Primers2: (Annealing Temperature 60.0 ºC)

F2: 5'-CTGTCCTGAGTGTGTGATGGTTC-3'
R1: 5'-AGGTCCTTTAACTCTACCCGGAA-3'
Product size: 541 bp

Homozygotes: one band with 610 bp
Heterozygotes: two bands with 610 bp and 541 bp
Wildtype allele: one band with 541 bp

**Figure S7. Generation of the *Arsa*-KO mouse model using CRISPR.**

### Supplemental Figure 7

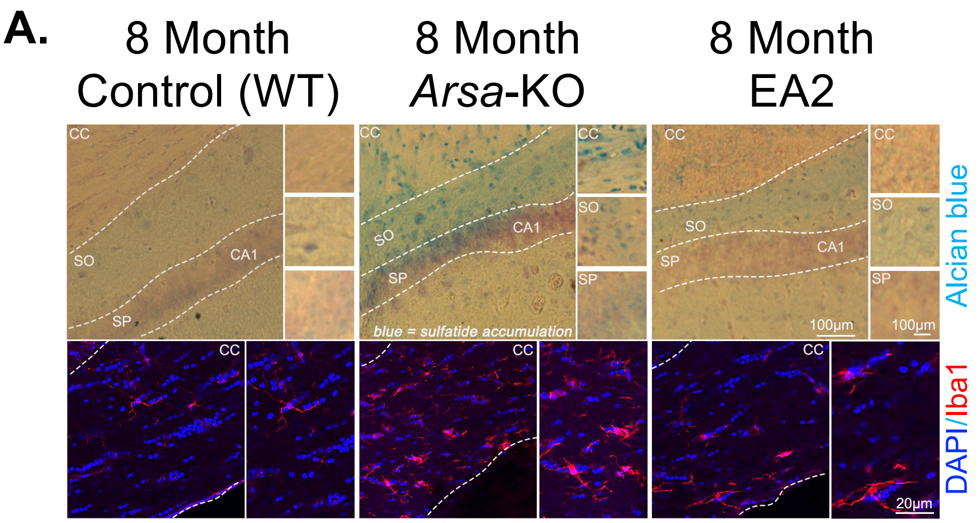

| Mouse | Chimerism % | VCN |
|---|---|---|
| EA2 R | 100 | 8.2 |
| EA2 L | 100 | 5.5 |
| EA2 RL | 100 | 3.8 |

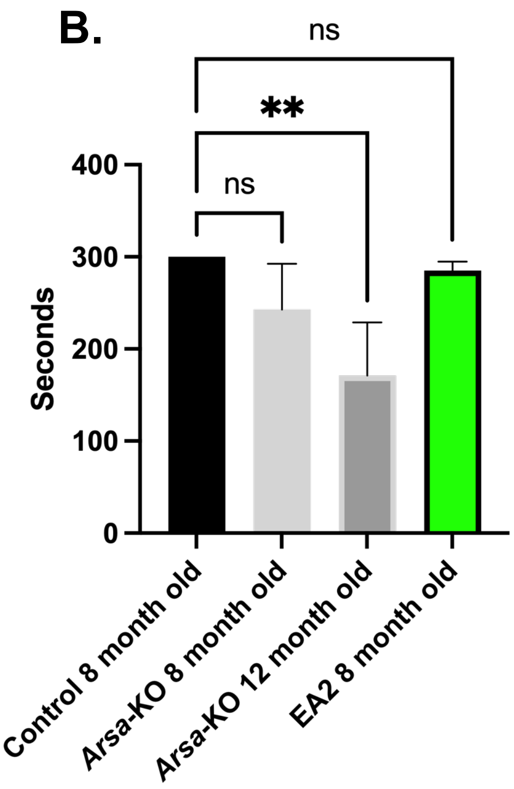

**Figure S8. Preliminary rotarod duration and pathological assessment with 8-12 months old *Arsa*-KO mice.** In pilot experiments, we assessed 8-month-old WT control mice, untreated *Arsa*-KO at 8 and 12 months old with 8-month-old high VCN EA2 transplanted mice to identify the ideal time range for detecting disease phenotype in the *Arsa*-KO mice and the efficacy of our LV system. (A) At 8 months, pathological brain sections showed already significant differences in sulfatide accumulation and microglia inflammation, indicating neural degeneration in the *Arsa*-KO mice. In EA2 treated mice, sulfatide accumulation and microglia inflammation was greatly reduced and similar to control mice, suggesting potent efficacy of our vector to ameliorate the disease. (B) Unlike 8-month-old *Arsa*-KO mice, 12-month-old *Arsa*-KO mice showed significant decline in duration on the rotarod compared to control and EA2 treated mice. EA2-treated mice demonstrated no significant differences in rotatord performance compared to control mice. **P<0.01.

**Supplemental Figure 8**

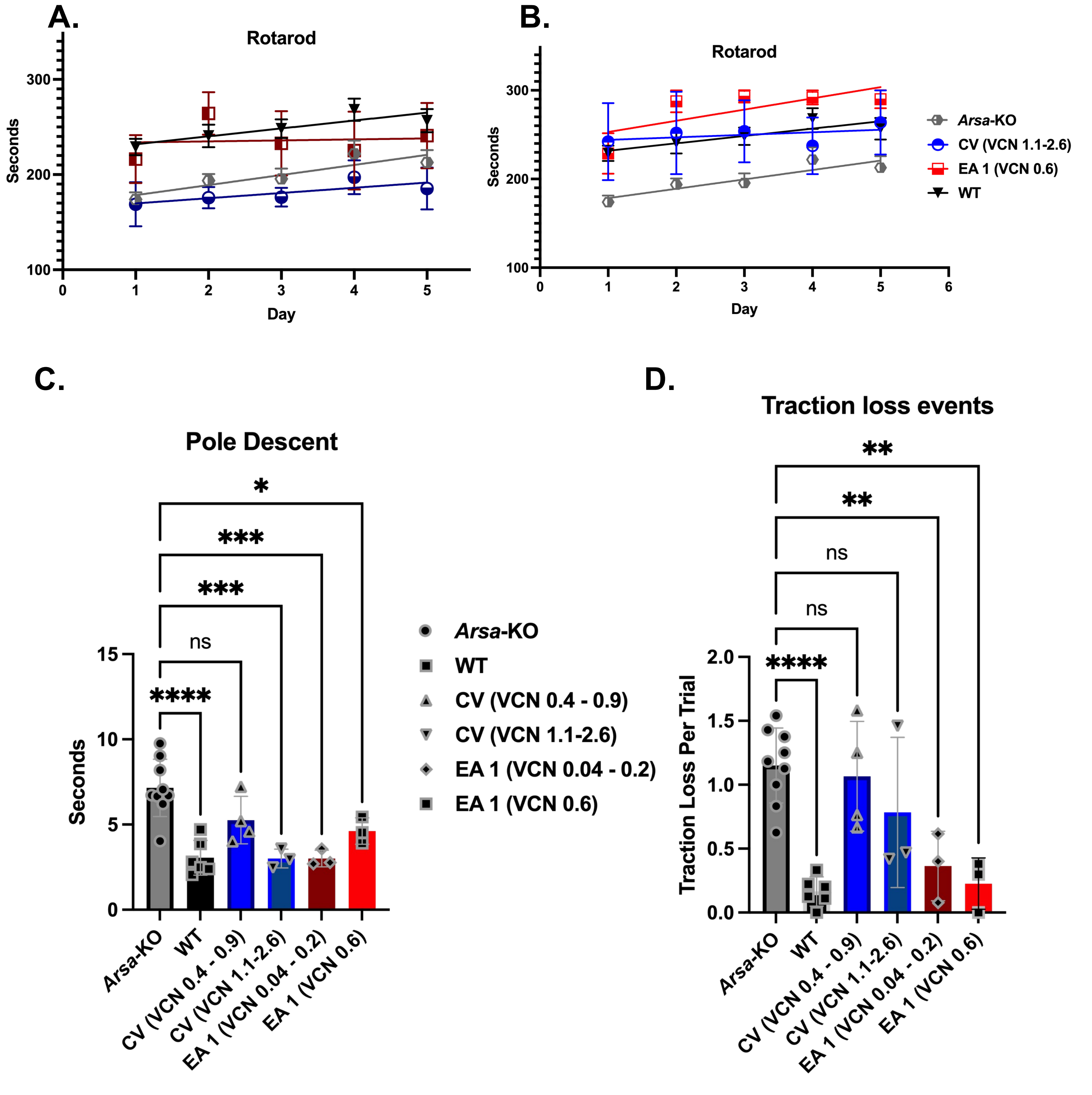

**Figure S9. Rotarod and pole descent motor control assays of untreated *Arsa*-KO disease mice compared to vector treated mice. (**A) Rotarod assessment over a 5-day time course for WT, untreated, CV or EA1 vector treated mice at lower VCN (CV 0.4-0.9 VCN and EA1 0.04-0.2) (n = range 3-9). (B) Rotarod assessment over a 5-day time course for WT, untreated, CV or EA1 vector treated mice at higher VCN (CV 1.1-2.6 VCN and EA1 0.6) (n = range 3-9). (C) Descent time duration for a predetermined segment of pole in vector treated groups compared to the *Arsa*-KO model (n = range 3-9). (D) Report of times mice slipped or lost grip during the pole descent assay (n = range 6-9). *P<0.05, **P<0.01, ***P<0.001, ****P<0.0001. Extended rotarod significance tables in supplement (**Supplemental Table S1** and **S2**).

#### Supplemental Figure 9

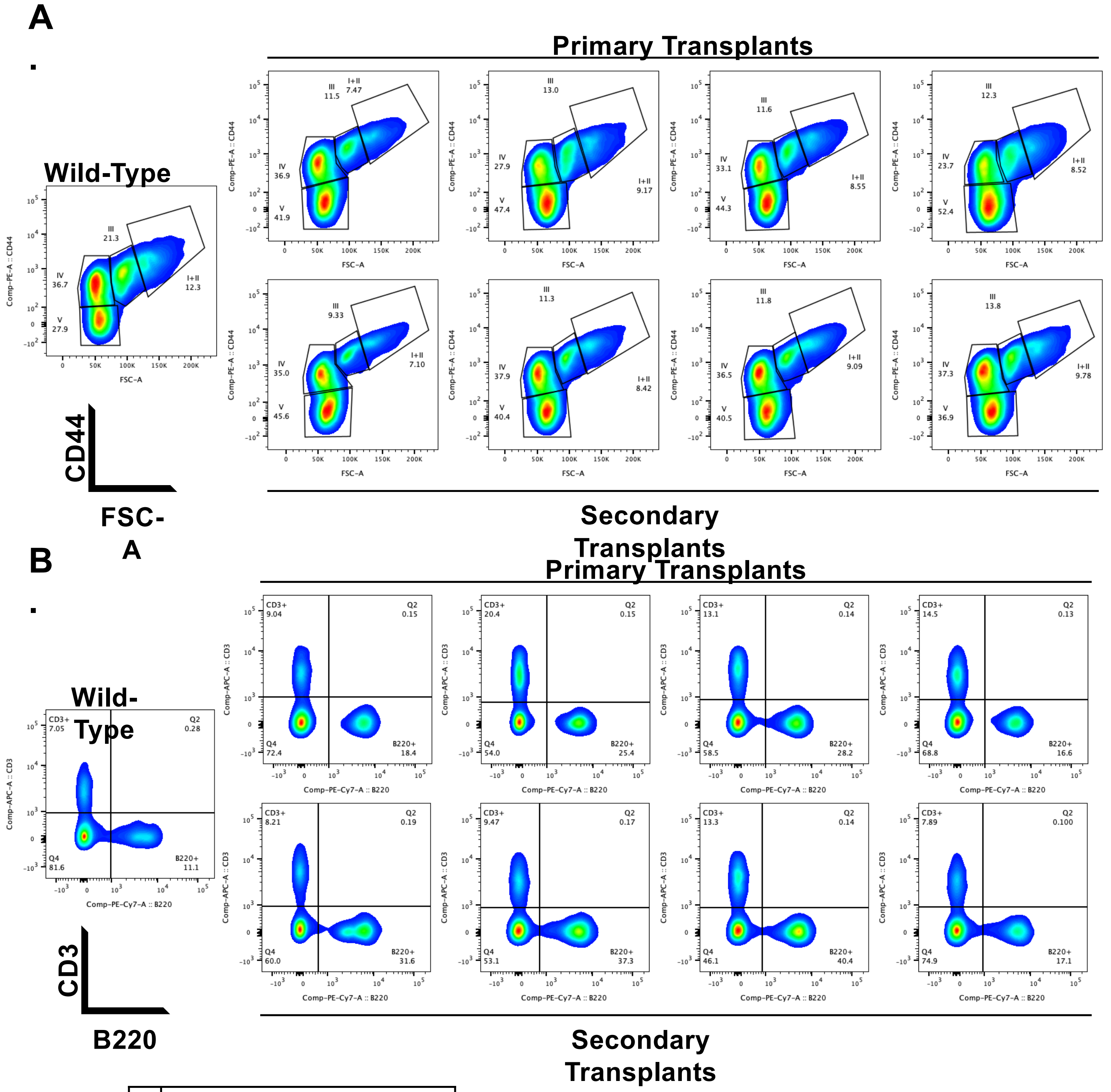

| | |
|---|---|
| I+II | Basophilic |
| III | Polychromatic |
| IV | Orthochromatic and Reticulocytes |
| V | Mature RBCs |

**Figure S10. *Arsa*-KO primary and secondary transplant flow cytometry analysis of bone marrow erythropoiesis and lymphopoiesis. (**A) Analysis of erythropoiesis (population I to V, from immature to enucleated red cells) by flow cytometry in 4 EA1 transduced primary (Top) and 4 secondary (Bottom) transplants. (B) Comparison of flow sort panel for lymphocyte B (B220) and T (CD3) cells in the same mice.

### Supplemental Figure 10

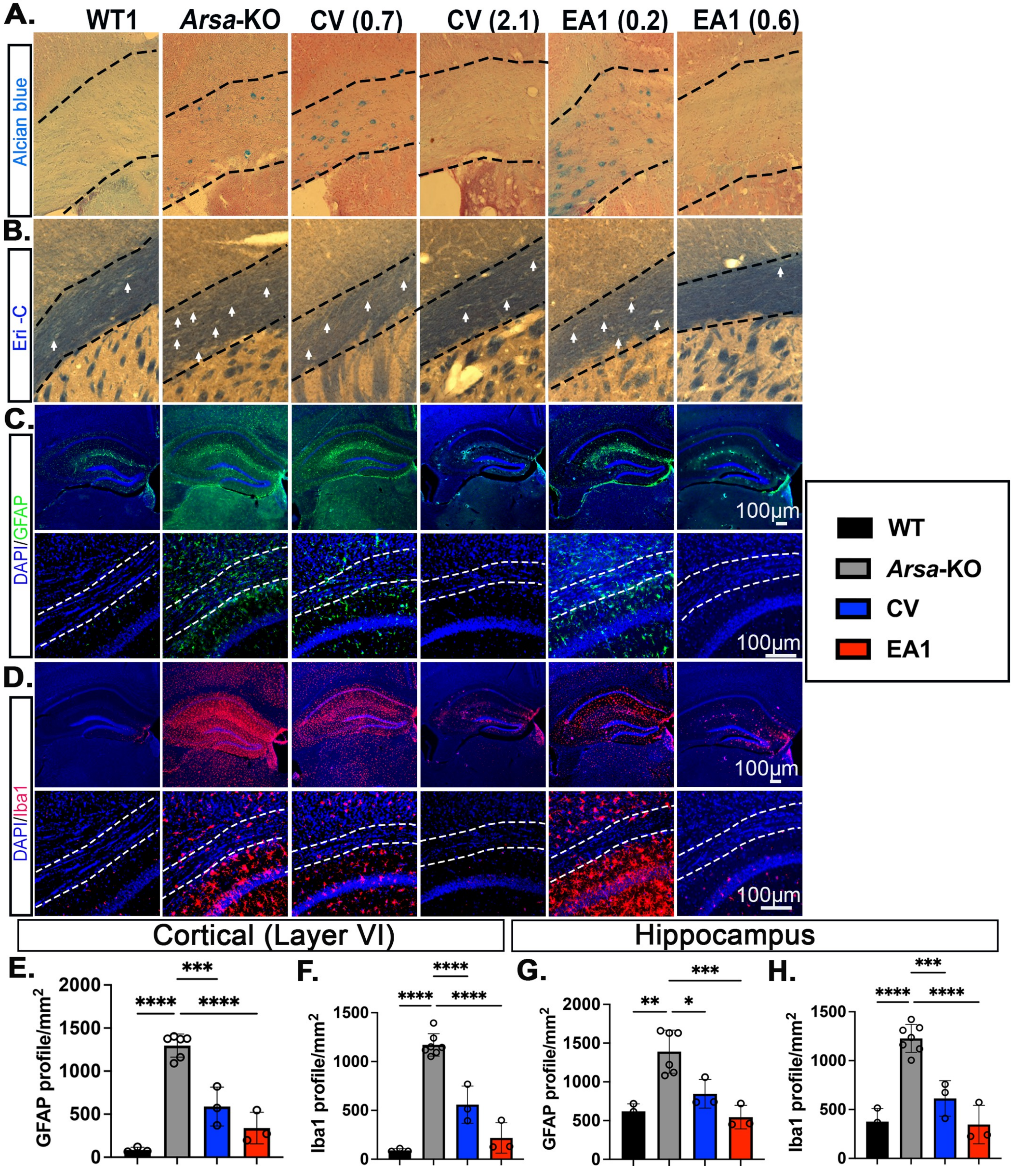

Supplemental Figure 11

**Figure S11. Neuropathological deficits of untreated *Arsa*-KO diseased mice relative to vector treated mice.** Treatment groups include WT, untreated, CV, and EA1 vectors treated at lower and higher VCN. (A) Representative images of sulfatide accumulation in the corpus callosum across treatment groups. (B) Representative myelin staining (Eri-C) images of the corpus callosum in treatment groups. (C) Representative immunofluorescent images of GFAP (astrocytes) in different brain areas. (D) Representative immunofluorescent images of Iba1 (microglia) in different brain areas. (E-H) Astrocytes (GFAP) and Microglia (Iba1) counts/mm$^2$ in cortex and hippocampus (layer shown in the image panel). *P<0.05, **P<0.01, ***P<0.001, ****P<0.0001. Extended significant tables in supplement **Table S3**.

### Legend Supplemental Figure 11

### Rotarod Low VCN Individual Day Stats

| | | | | | |
|---|---|---|---|---|---|
| Within each row, compare columns (simple effects within rows) | | | | | |
| Number of families | 5 | | | | |
| Number of comparisons per family | 6 | | | | |
| Alpha | 0.05 | | | | |
| Tukey's multiple comparisons test | Mean Diff. | 95.00% CI of diff. | Below threshold? | Summary | Adjusted P Value |
| Day 1 | | | | | |
| *Arsa*-KO vs. EA1 (VCN 0.04 - 0.2) | -42.30 | -196.4 to 111.8 | No | ns | 0.5074 |
| *Arsa*-KO vs. CV (VCN 0.4 - 0.9) | 5.176 | -99.45 to 109.8 | No | ns | 0.9960 |
| *Arsa*-KO vs. WT | -55.03 | -88.26 to -21.79 | Yes | ** | 0.0016 |
| EA1 (VCN 0.04 - 0.2) vs. CV (VCN 0.4 - 0.9) | 47.47 | -83.04 to 178.0 | No | ns | 0.5600 |
| EA1 (VCN 0.04 - 0.2) vs. WT | -12.73 | -161.6 to 136.1 | No | ns | 0.9582 |
| CV (VCN 0.4 - 0.9) vs. WT | -60.20 | -163.1 to 42.72 | No | ns | 0.2172 |
| Day 2 | | | | | |
| *Arsa*-KO vs. EA1 (VCN 0.04 - 0.2) | -70.65 | -206.1 to 64.83 | No | ns | 0.1926 |
| *Arsa*-KO vs. CV (VCN 0.4 - 0.9) | 17.94 | -29.31 to 65.18 | No | ns | 0.5648 |
| *Arsa*-KO vs. WT | -46.93 | -88.63 to -5.234 | Yes | * | 0.0272 |
| EA1 (VCN 0.04 - 0.2) vs. CV (VCN 0.4 - 0.9) | 88.58 | -31.98 to 209.1 | No | ns | 0.1093 |
| EA1 (VCN 0.04 - 0.2) vs. WT | 23.71 | -93.60 to 141.0 | No | ns | 0.7900 |
| CV (VCN 0.4 - 0.9) vs. WT | -64.87 | -116.6 to -13.16 | Yes | * | 0.0160 |
| Day 3 | | | | | |
| *Arsa*-KO vs. EA1 (VCN 0.04 - 0.2) | -37.37 | -240.3 to 165.6 | No | ns | 0.7406 |
| *Arsa*-KO vs. CV (VCN 0.4 - 0.9) | 19.27 | -25.56 to 64.10 | No | ns | 0.5682 |
| *Arsa*-KO vs. WT | -52.67 | -95.11 to -10.23 | Yes | * | 0.0135 |
| EA1 (VCN 0.04 - 0.2) vs. CV (VCN 0.4 - 0.9) | 56.64 | -151.2 to 264.5 | No | ns | 0.5088 |
| EA1 (VCN 0.04 - 0.2) vs. WT | -15.30 | -222.1 to 191.5 | No | ns | 0.9675 |
| CV (VCN 0.4 - 0.9) vs. WT | -71.94 | -116.3 to -27.60 | Yes | ** | 0.0036 |
| Day 4 | | | | | |
| *Arsa*-KO vs. EA1 (VCN 0.04 - 0.2) | -3.333 | -245.5 to 238.8 | No | ns | 0.9998 |
| *Arsa*-KO vs. CV (VCN 0.4 - 0.9) | 24.72 | -50.35 to 99.80 | No | ns | 0.7024 |
| *Arsa*-KO vs. WT | -46.21 | -99.00 to 6.589 | No | ns | 0.0958 |
| EA1 (VCN 0.04 - 0.2) vs. CV (VCN 0.4 - 0.9) | 28.06 | -200.9 to 257.0 | No | ns | 0.9161 |
| EA1 (VCN 0.04 - 0.2) vs. WT | -42.87 | -294.3 to 208.6 | No | ns | 0.7624 |
| CV (VCN 0.4 - 0.9) vs. WT | -70.93 | -146.0 to 4.174 | No | ns | 0.0622 |
| Day 5 | | | | | |
| *Arsa*-KO vs. EA1 (VCN 0.04 - 0.2) | -28.22 | -225.0 to 168.6 | No | ns | 0.8638 |
| *Arsa*-KO vs. CV (VCN 0.4 - 0.9) | 27.36 | -65.35 to 120.1 | No | ns | 0.7197 |
| *Arsa*-KO vs. WT | -43.89 | -95.84 to 8.058 | No | ns | 0.1115 |
| EA1 (VCN 0.04 - 0.2) vs. CV (VCN 0.4 - 0.9) | 55.58 | -120.1 to 231.3 | No | ns | 0.5813 |
| EA1 (VCN 0.04 - 0.2) vs. WT | -15.67 | -215.9 to 184.6 | No | ns | 0.9686 |
| CV (VCN 0.4 - 0.9) vs. WT | -71.25 | -164.8 to 22.28 | No | ns | 0.1235 |

**Table S1. Table of statical significance for every individual day of rotarod, 1-5, in the low VCN group on rotarod in figure S10 A.** Statistical significance for comparison between each *Arsa*-KO treatment group of low VCN CV and EA1 in addition to WT and *Arsa*-KO mice.

#### Supplemental Table 1

### Rotarod High VCN Individual Day Stats

| | | | | | | |
|---|---|---|---|---|---|---|
| | Within each row, compare columns (simple effects within rows) | | | | | |
| | Number of families | 5 | | | | |
| | Number of comparisons per family | 10 | | | | |
| | Alpha | 0.05 | | | | |
| | Tukey's multiple comparisons test | Mean Diff. | 95.00% CI of diff. | Below threshold? | Summary | Adjusted P Value |
| Day 1 | *Arsa*-KO vs. EA2 | 17.81 | -23.91 to 59.54 | No | ns | 0.6353 |
| | *Arsa*-KO vs. EA1 (VCN 0.6) | -54.85 | -206.5 to 96.81 | No | ns | 0.3742 |
| | *Arsa*-KO vs. CV (VCN 1.1-2.6) | -68.30 | -387.3 to 250.7 | No | ns | 0.6219 |
| | *Arsa*-KO vs. WT | -55.03 | -90.68 to -19.37 | Yes | ** | 0.0024 |
| | EA2 vs. EA1 (VCN 0.6) | -72.67 | -211.2 to 65.90 | No | ns | 0.2260 |
| | EA2 vs. CV (VCN 1.1-2.6) | -86.11 | -392.3 to 220.1 | No | ns | 0.4855 |
| | EA2 vs. WT | -72.84 | -116.7 to -28.94 | Yes | ** | 0.0019 |
| | EA1 (VCN 0.6) vs. CV (VCN 1.1-2.6) | -13.44 | -272.4 to 245.5 | No | ns | 0.9981 |
| | EA1 (VCN 0.6) vs. WT | -0.1746 | -145.9 to 145.5 | No | ns | >0.9999 |
| | CV (VCN 1.1-2.6) vs. WT | 13.27 | -300.5 to 327.0 | No | ns | 0.9969 |
| Day2 | *Arsa*-KO vs. EA2 | 24.57 | -37.53 to 86.67 | No | ns | 0.6345 |
| | *Arsa*-KO vs. EA1 (VCN 0.6) | -93.98 | -163.5 to -24.44 | Yes | * | 0.0203 |
| | *Arsa*-KO vs. CV (VCN 1.1-2.6) | -58.20 | -402.5 to 286.1 | No | ns | 0.7453 |
| | *Arsa*-KO vs. WT | -46.93 | -91.80 to -2.070 | Yes | * | 0.0395 |
| | EA2 vs. EA 1 (VCN 0.6) | -118.6 | -191.3 to -45.84 | Yes | ** | 0.0043 |
| | EA2 vs. CV (VCN 1.1-2.6) | -82.78 | -386.2 to 220.6 | No | ns | 0.5582 |
| | EA2 vs. WT | -71.51 | -137.0 to -5.990 | Yes | * | 0.0316 |
| | EA1 (VCN 0.6) vs. CV (VCN 1.1-2.6) | 35.78 | -285.6 to 357.2 | No | ns | 0.9287 |
| | EA1 (VCN 0.6) vs. WT | 47.05 | -17.95 to 112.0 | No | ns | 0.1592 |
| | CV (VCN 1.1-2.6) vs. WT | 11.27 | -311.9 to 334.4 | No | ns | 0.9988 |
| Day 3 | *Arsa*-KO vs. EA2 | 2.185 | -63.62 to 67.99 | No | ns | >0.9999 |
| | *Arsa*-KO vs. EA1 (VCN 0.6) | -97.59 | -140.0 to -55.16 | Yes | *** | 0.0002 |
| | *Arsa*-KO vs. CV (VCN 1.1-2.6) | -58.26 | -292.9 to 176.4 | No | ns | 0.5985 |
| | *Arsa*-KO vs. WT | -52.67 | -98.17 to -7.172 | Yes | * | 0.0202 |
| | EA2 vs. EA1 (VCN 0.6) | -99.78 | -165.2 to -34.35 | Yes | ** | 0.0065 |
| | EA2 vs. CV (VCN 1.1-2.6) | -60.44 | -270.0 to 149.1 | No | ns | 0.5978 |
| | EA2 vs. WT | -54.86 | -120.4 to 10.71 | No | ns | 0.1093 |
| | EA1 (VCN 0.6) vs. CV (VCN 1.1-2.6) | 39.33 | -214.0 to 292.7 | No | ns | 0.8015 |
| | EA1 (VCN 0.6) vs. WT | 44.92 | 2.994 to 86.85 | Yes | * | 0.0360 |
| | CV (VCN 1.1-2.6) vs. WT | 5.587 | -233.5 to 244.7 | No | ns | 0.9998 |
| Day 4 | *Arsa*-KO vs. EA2 | 16.11 | -45.77 to 77.99 | No | ns | 0.9176 |
| | *Arsa*-KO vs. EA1 (VCN 0.6) | -70.11 | -123.0 to -17.27 | Yes | ** | 0.0097 |
| | *Arsa*-KO vs. CV (VCN 1.1-2.6) | -15.56 | -209.7 to 178.6 | No | ns | 0.9875 |
| | *Arsa*-KO vs. WT | -46.21 | -102.8 to 10.39 | No | ns | 0.1360 |
| | EA2 vs. EA1 (VCN 0.6) | -86.22 | -143.1 to -29.32 | Yes | ** | 0.0061 |
| | EA2 vs. CV (VCN 1.1-2.6) | -31.67 | -227.2 to 163.9 | No | ns | 0.8761 |
| | EA2 vs. WT | -62.32 | -121.3 to -3.315 | Yes | * | 0.0374 |
| | EA1 (VCN 0.6) vs. CV (VCN 1.1-2.6) | 54.56 | -168.3 to 277.4 | No | ns | 0.5760 |
| | EA1 (VCN 0.6) vs. WT | 23.90 | -25.44 to 73.25 | No | ns | 0.4929 |
| | CV (VCN 1.1-2.6) vs. WT | -30.65 | -234.8 to 173.5 | No | ns | 0.8791 |
| Day 5 | *Arsa*-KO vs. EA2 | -13.17 | -95.43 to 69.09 | No | ns | 0.9805 |
| | *Arsa*-KO vs. EA1 (VCN 0.6) | -77.11 | -133.4 to -20.80 | Yes | ** | 0.0091 |
| | *Arsa*-KO vs. CV (VCN 1.1-2.6) | -51.00 | -284.1 to 182.1 | No | ns | 0.7048 |
| | *Arsa*-KO vs. WT | -43.89 | -99.58 to 11.80 | No | ns | 0.1571 |
| | EA2 vs. EA1 (VCN 0.6) | -63.94 | -147.1 to 19.26 | No | ns | 0.1399 |
| | EA2 vs. CV (VCN 1.1-2.6) | -37.83 | -242.4 to 166.7 | No | ns | 0.8795 |
| | EA2 vs. WT | -30.72 | -112.9 to 51.49 | No | ns | 0.7092 |
| | EA1 (VCN 0.6) vs. CV (VCN 1.1-2.6) | 26.11 | -222.3 to 274.5 | No | ns | 0.9424 |
| | EA1 (VCN 0.6) vs. WT | 33.22 | -23.38 to 89.83 | No | ns | 0.3171 |
| | CV (VCN 1.1-2.6) vs. WT | 7.111 | -230.0 to 244.2 | No | ns | 0.9995 |

**Table S2. Table of statical significance for every individual day, 1-5, in the high VCN group on rotarod in figure S10 B.** Statistical significance for comparison between each Arsa-KO treatment group of high VCN CV and EA1 in addition to WT and Arsa-KO mice.

#### Supplemental Table 2

## A.

| Sulfatide accumulation (in corpus callosum) | | | | |
|---|---|---|---|---|
| | WT control | *Arsa*-KO | CV (0.4-0.6) | EA1 (0.6) |
| Number of values | 3 | 6 | 4 | 3 |
| Minimum | 20 | 433 | 145.3 | 14.95 |
| Maximum | 69 | 805 | 418 | 167.7 |
| Range | 49 | 372 | 272.8 | 152.7 |
| Mean | 52.33 | 634 | 255.6 | 76.71 |
| Std. Deviation | 28.01 | 130 | 125.4 | 80.43 |
| Std. Error of Mean | 16.17 | 53.09 | 62.7 | 46.43 |

## B.

| Eri-C staining (in corpus callosum) | | | | |
|---|---|---|---|---|
| | WT control | *Arsa*-KO | CV (0.4-0.6) | EA1 (0.6) |
| Number of values | 3 | 7 | 4 | 3 |
| Minimum | 11 | 17 | 15 | 9 |
| Maximum | 12 | 47 | 23 | 12 |
| Range | 1 | 30 | 8 | 3 |
| Mean | 11.67 | 26.77 | 18.5 | 10.67 |
| Std. Deviation | 0.5774 | 10.35 | 3.416 | 1.528 |
| Std. Error of Mean | 0.3333 | 3.91 | 1.708 | 0.8819 |

**Table S3: Counts and descriptive statistics of histological section from untreated and treated *Arsa*-KO mice from Figure 7A**, staining for sulfatide, and **Figure 7B**, staining for Eri-C.

**Supplemental Table 3**

C.

| GFAP staining( in corpus callosum) | | | | |
|---|---|---|---|---|
| | WT control | *Arsa*-KO | CV (0.6-0.9) | EA1 (0.6) |
| Number of values | 3 | 6 | 3 | 3 |
| Minimum | 272 | 782 | 696 | 457 |
| Maximum | 302 | 2008 | 1188 | 742 |
| Range | 30 | 1226 | 492 | 285 |
| Mean | 290 | 1480 | 923.3 | 558.2 |
| Std. Deviation | 15.87 | 404.2 | 248.1 | 159.4 |
| Std. Error of Mean | 9.165 | 165 | 143.2 | 92.04 |
| **GFAP staining (in Cortex)** | | | | |
| | WT control | *Arsa*-KO | CV (0.6-0.9) | EA1 (0.6) |
| Number of values | 3 | 6 | 3 | 3 |
| Minimum | 64 | 1091 | 370 | 198 |
| Maximum | 120 | 1402 | 822 | 547 |
| Range | 56 | 311 | 452 | 349 |
| Mean | 85 | 1296 | 588.3 | 339 |
| Std. Deviation | 30.51 | 134.3 | 226.4 | 183.9 |
| Std. Error of Mean | 17.62 | 54.83 | 130.7 | 106.2 |
| **GFAP staining (in hippocampus)** | | | | |
| | WT control | *Arsa*-KO | CV (0.6-0.9) | EA1 (0.6) |
| Number of values | 3 | 6 | 3 | 3 |
| Minimum | 526 | 1082 | 738 | 450 |
| Maximum | 718 | 1659 | 1060 | 721 |
| Range | 192 | 577 | 322 | 271 |
| Mean | 619.3 | 1390 | 846.3 | 544.7 |
| Std. Deviation | 96.11 | 277.2 | 185 | 152.8 |
| Std. Error of Mean | 55.49 | 113.2 | 106.8 | 88.25 |

**Table S3: Counts and descriptive statistics of histological section from untreated and treated *Arsa*-KO mice from Figure 7C**, staining for GFAP.

**Supplemental Table 3**

**D.**

| Iba1 staining (in corpus callosum) | | | | |
|---|---|---|---|---|
| | WT | *Arsa*-KO | CV (0.4-0.6) | EA1 (0.6) |
| Number of values | 3 | 7 | 3 | 3 |
| Minimum | 96 | 716 | 355.4 | 128 |
| Maximum | 212 | 964 | 546.2 | 384.4 |
| Range | 116 | 248 | 190.8 | 256.4 |
| Mean | 148.7 | 778.7 | 447.2 | 246.5 |
| Std. Deviation | 58.73 | 88 | 95.59 | 129.3 |
| Std. Error of Mean | 33.91 | 33.26 | 55.19 | 74.66 |
| Iba1 staining (in cortex) | | | | |
| | WT | *Arsa*-KO | CV (0.4-0.6) | EA1 (0.6) |
| Number of values | 3 | 7 | 3 | 3 |
| Minimum | 81 | 1052 | 394.6 | 126 |
| Maximum | 110 | 1394 | 768.5 | 398.9 |
| Range | 29 | 342.7 | 373.8 | 272.9 |
| Mean | 91 | 1170 | 559 | 218.3 |
| Std. Deviation | 16.46 | 114.7 | 191 | 156.4 |
| Std. Error of Mean | 9.504 | 43.35 | 110.3 | 90.3 |
| Iba1 staining (in hippocampus) | | | | |
| | WT | *Arsa*-KO | CV (0.4-0.6) | EA1 (0.6) |
| Number of values | 3 | 7 | 3 | 3 |
| Minimum | 249 | 1004 | 409 | 225 |
| Maximum | 515 | 1422 | 758 | 573.3 |
| Range | 266 | 418 | 349 | 348.3 |
| Mean | 377.3 | 1227 | 613.7 | 346.4 |
| Std. Deviation | 133.2 | 142.9 | 182.2 | 196.7 |
| Std. Error of Mean | 76.93 | 54.03 | 105.2 | 113.5 |

**Table S3: Counts and descriptive statistics of histological section from untreated and treated *Arsa*-KO mice from Figure 7D**, staining for Iba1.

#### Supplemental Table 3

| Status | Guide Target | PAM Sequence | Indel % | Model Fit ($R^2$) | Knockout-Score |
|---|---|---|---|---|---|
| Succeeded | CGGGAGCCGGCCGGTCAGGA | GGG | 77 | 0.96 | 72 |

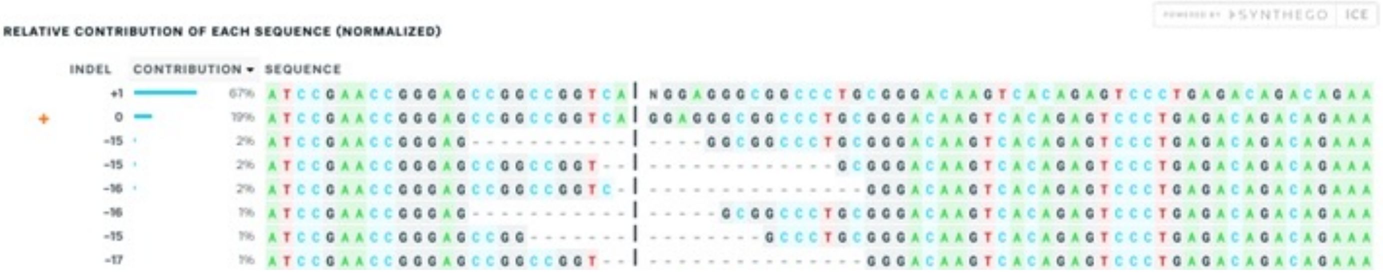

The contributions show the inferred sequences present in your edited population and their relative proportions (in contrast to the Indel plot (Indel Distribution tab) that does not specify sequence contributions). Cut sites are represented by black vertical dotted lines, and the wild-type sequence is marked by a "+" symbol on the far left.

| Status | Guide Target | PAM Sequence | Indel % | Model Fit ($R^2$) | Knockout-Score |
|---|---|---|---|---|---|
| Succeeded | CGGGAGCCGGCCGGTCAGGA | GGG | 77 | 0.96 | 72 |

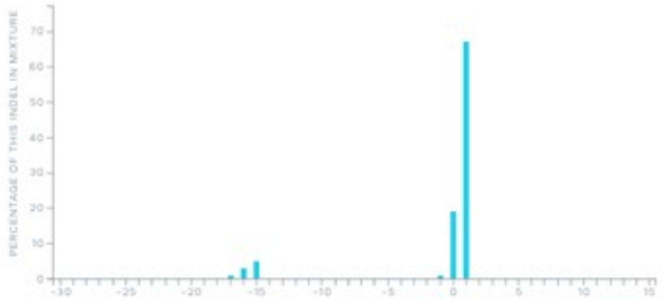

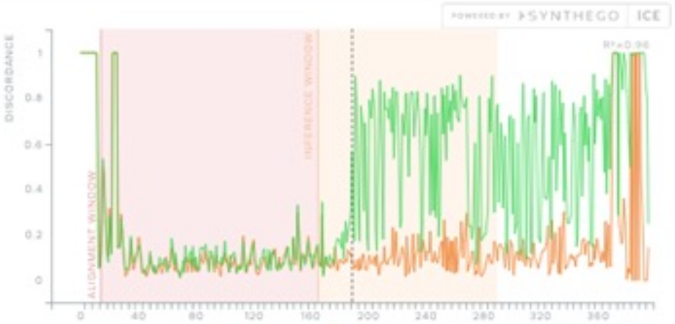

The Indel plot displays the inferred distribution of indels in the entire edited population of genomes. Hovering over each bar of the Indel plot shows the size of the insertion or deletion (+ or - 1 or more nucleotides), along with the percentage of genomes that contain it. Note: Each indel size represented in the Indel plot may not necessarily occur in the same sequence. The percentages of different indel sizes in the cell population are not the same as ICE/KO-Score scores.

The discordance plot details the level of alignment per base between the wild type (control) and the edited sample in the inference window (the region around the cute site), i.e. it shows the average amount of signal that disagrees with the reference sequence derived from the control trace file. On the plot, the green line and orange line should be close together before the cut site, with a typical CRISPR edit resulting in a jump in the discordance near the cutsite and continuing to remain far apart after the cut site (representing a high level of sequence discordance).

| Status | Guide Target | PAM Sequence | Indel % | Model Fit ($R^2$) | Knockout-Score |
|---|---|---|---|---|---|
| Succeeded | CGGGAGCCGGCCGGTCAGGA | GGG | 77 | 0.96 | 72 |

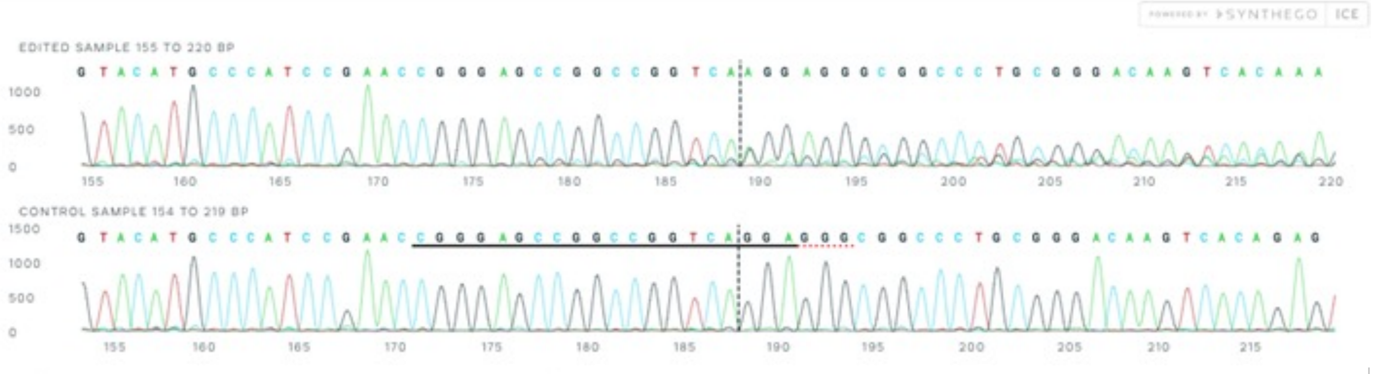

This is the Sanger sequence view showing edited and wild-type (control) sequences in the region around the guide sequence. This shows sequence base calls from both the control and the experimental sample .ab1 files, which will contain mixed base calls. The horizontal black underlined region represents the guide sequence. The horizontal red underline is the PAM site. The vertical black dotted line represents the actual cut site. Cutting and error-prone repair usually results in mixed sequencing bases after the cut.

#### Supplemental Table 4

| Tube | Label | Guide Sequence | # Cells / Yield |
|---|---|---|---|
| 1 | ARSA+50627396 Knockout – HMC3 p3 | CGGGAGCCGGCCGGUCAGGA | $10^6$ cells |
| 2 | ARSA+50627396 Knockout – HMC3 p3 | CGGGAGCCGGCCGGUCAGGA | $10^6$ cells |
| 3 | HMC3 WT p3 | None | $10^6$ cells |
| 4 | HMC3 WT p3 | None | $10^6$ cells |

**Table S4. Generation of the HMC3 *ARSA-KO* cell line using CRISPR.**

### Supplemental Table 4

| ddPCR | Target Gene: | cDNA /gDNA | Oligo type | Fluorophore on probe | Sequence |
|---|---|---|---|---|---|
| VCN in human cells | Psi | gDNA | Primer-F: | | ACCTGAAAGCGAAAGGGAAAC |
| | | | Primer-R: | | CGCACCCATCTCTCTCCTTCT |
| | | | Probe: | FAM | AGCTCTCTCGACGCAGGACTC GGC |
| | RPP30 | gDNA | Primer-F: | | ACCACCACATCCCAGCTAAT |
| | | | Primer-R: | | GACTGCTTGAATCTGCCAGG |
| | | | Probe: | HEX | TGCCATGTCCAGGCTAGTCTCA AA |
| mouse femal/male chimerism | mZfy1 | gDNA | Primer-F: | | AAGTGCTTCTTGGCATCACC |
| | | | Primer-R: | | ATCCAAAGACTGCTCCACCT |
| | | | Probe: | FAM | TGGGTTTGGTGTCTGCAGATG GT |
| | PCBP2 | gDNA | Primer-F: | | CCAGTCTGCTTGGCATGAAA |
| | | | Primer-R: | | GGTACCCTTAGCAGCAGACA |
| | | | Probe: | HEX | CCCATCCCTCTCCTGGCTCTAA |

**Table S5. Table of primer sequences used for ddPCR.** Primer sequences used for VCN determination in human and mouse cells and to determine BMT chimerism (male bone marrow into female recipient or vice versa).

**Supplemental Table 5**

| CV | uL used | ml used | # of HSC | Viral Titer | MOI |
|---|---|---|---|---|---|
| 3/17/22 | 0.3 | 0.0003 | 1.00E+06 | 2.2667E+10 | 6.8 |
| 3/17/22 | 0.3 | 0.0003 | 1.00E+06 | 2.2667E+10 | 6.8 |
| 5/13/22 | 0.05 | 0.00005 | 9.00E+05 | 2.2667E+10 | 1.26 |
| 5/13/22 | 0.05 | 0.00005 | 9.00E+05 | 2.2667E+10 | 1.26 |
| 5/13/22 | 0.05 | 0.00005 | 9.00E+05 | 2.2667E+10 | 1.26 |
| 5/13/22 | 0.05 | 0.00005 | 9.00E+05 | 2.2667E+10 | 1.26 |
| 5/13/22 | 0.05 | 0.00005 | 9.00E+05 | 2.2667E+10 | 1.26 |
| **EA1** | **uL used** | **ml used** | **# of HSC** | **Viral Titer** | **MOI** |
| 3/17/22 | 0.55 | 0.00055 | 1.00E+06 | 1.08E+09 | 0.59 |
| 3/17/22 | 0.55 | 0.00055 | 1.00E+06 | 1.08E+09 | 0.59 |
| 2/17/23 | 1.1 | 0.0011 | 1.33E+06 | 4.00E+08 | 0.33 |
| 2/17/23 | 1.1 | 0.0011 | 1.33E+06 | 4.00E+08 | 0.33 |
| 2/17/23 | 1.1 | 0.0011 | 1.33E+06 | 4.00E+08 | 0.33 |
| 2/17/23 | 1.1 | 0.0011 | 1.33E+06 | 4.00E+08 | 0.33 |
| 2/17/23 | 1.1 | 0.0011 | 1.33E+06 | 4.00E+08 | 0.33 |
| **EA2** | **uL used** | **ml used** | **# of HSC** | **Viral Titer** | **MOI** |
| 1/25/22 | 1 | 0.001 | 1.00E+06 | 4.60E+09 | 4.6 |
| 1/25/22 | 1 | 0.001 | 1.00E+06 | 4.60E+09 | 4.6 |
| 1/25/22 | 1 | 0.001 | 1.00E+06 | 4.60E+09 | 4.6 |

**Table S6. Table of viral titers and MOI from baches of virus used for *Arsa*-KO lentiviral treated mice.** MOI, cell counts and titers for CV, EA1 and EA2-treated mice.

### Supplemental Table 6

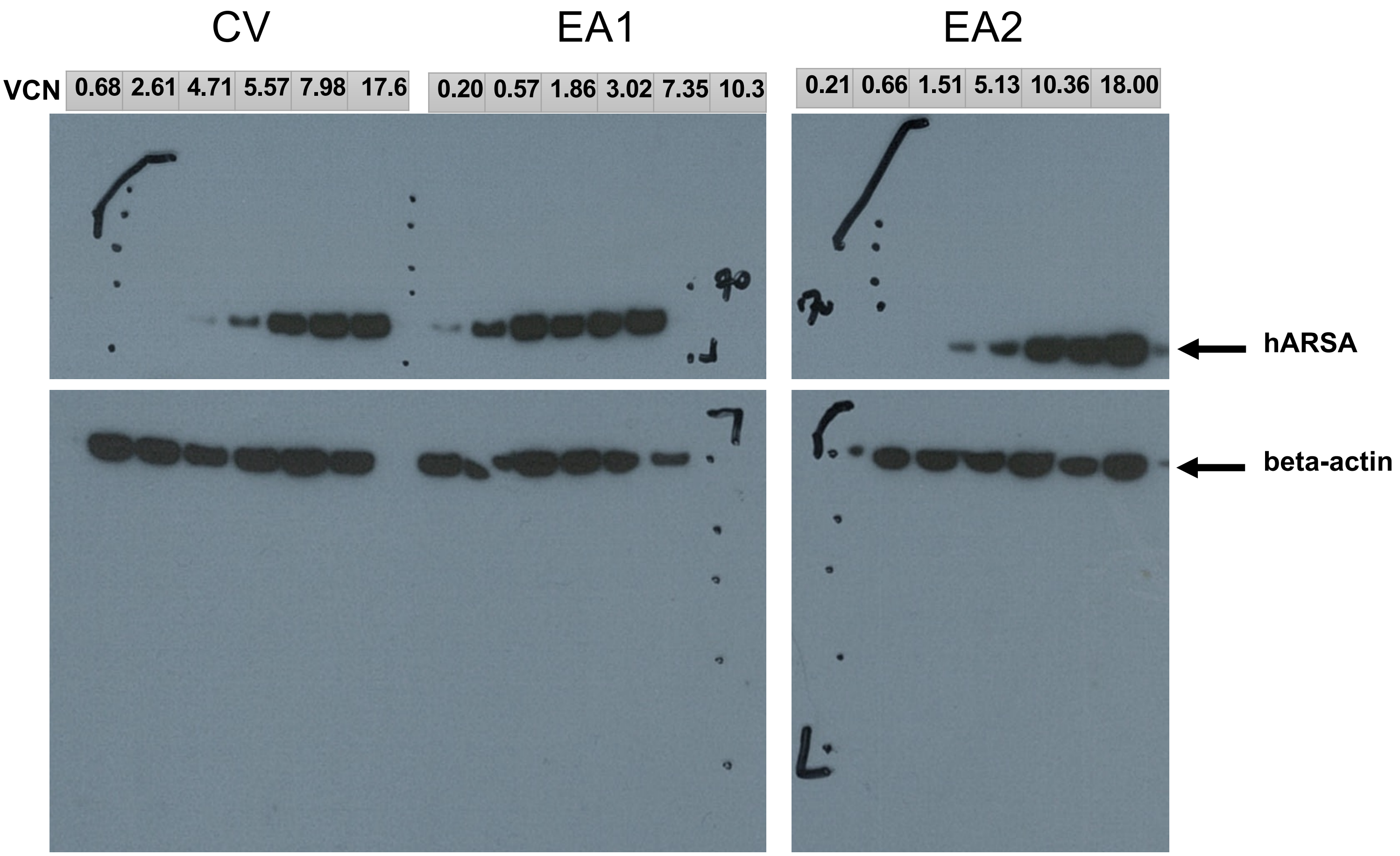

**Table S7**. Raw Fibroblast audioradiogram from Figure 2C.

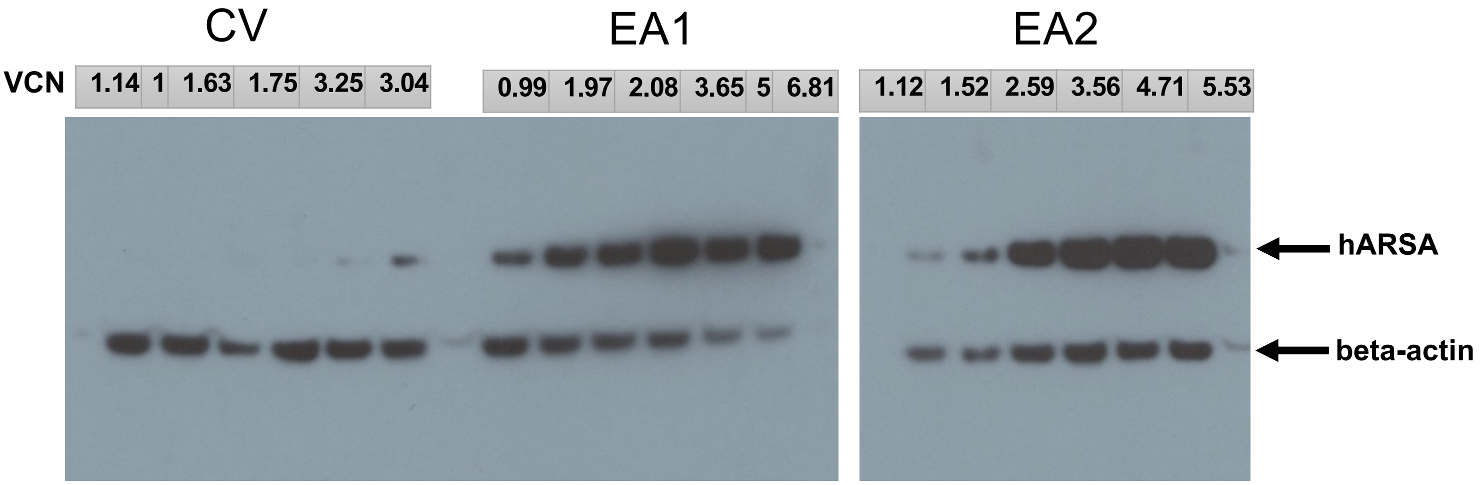

**Table S8**. Raw HMC3 audioradiogram from Figure 2D.

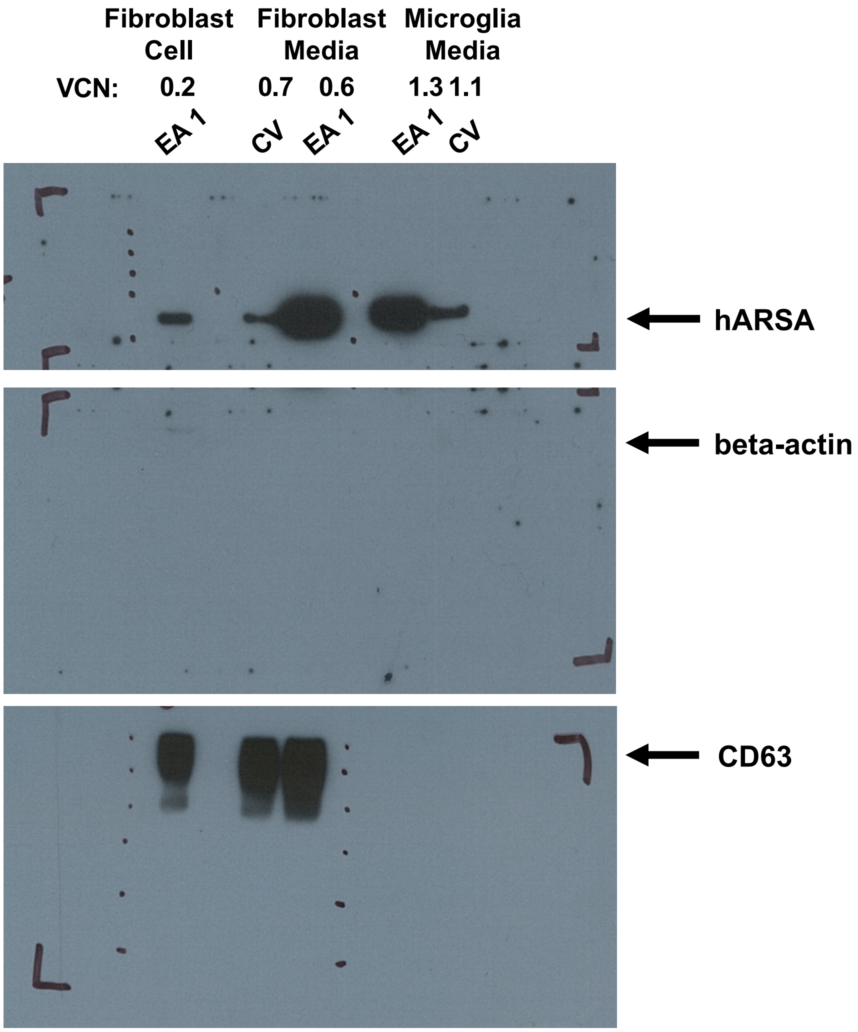

**Table S9**. Raw audioradiogram from Figure 3A.

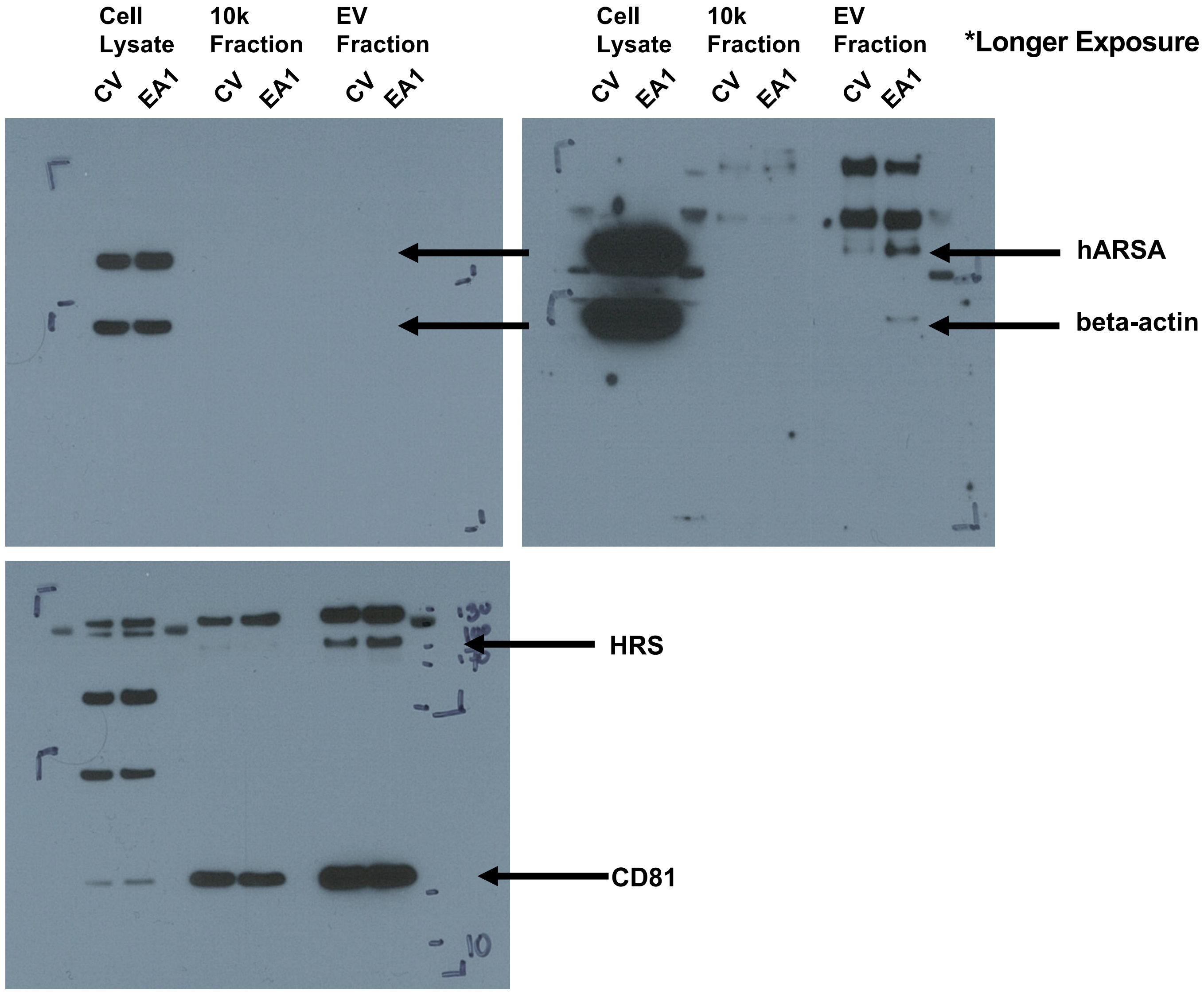

**Table S10**. Raw audioradiogram from Figure 3B.

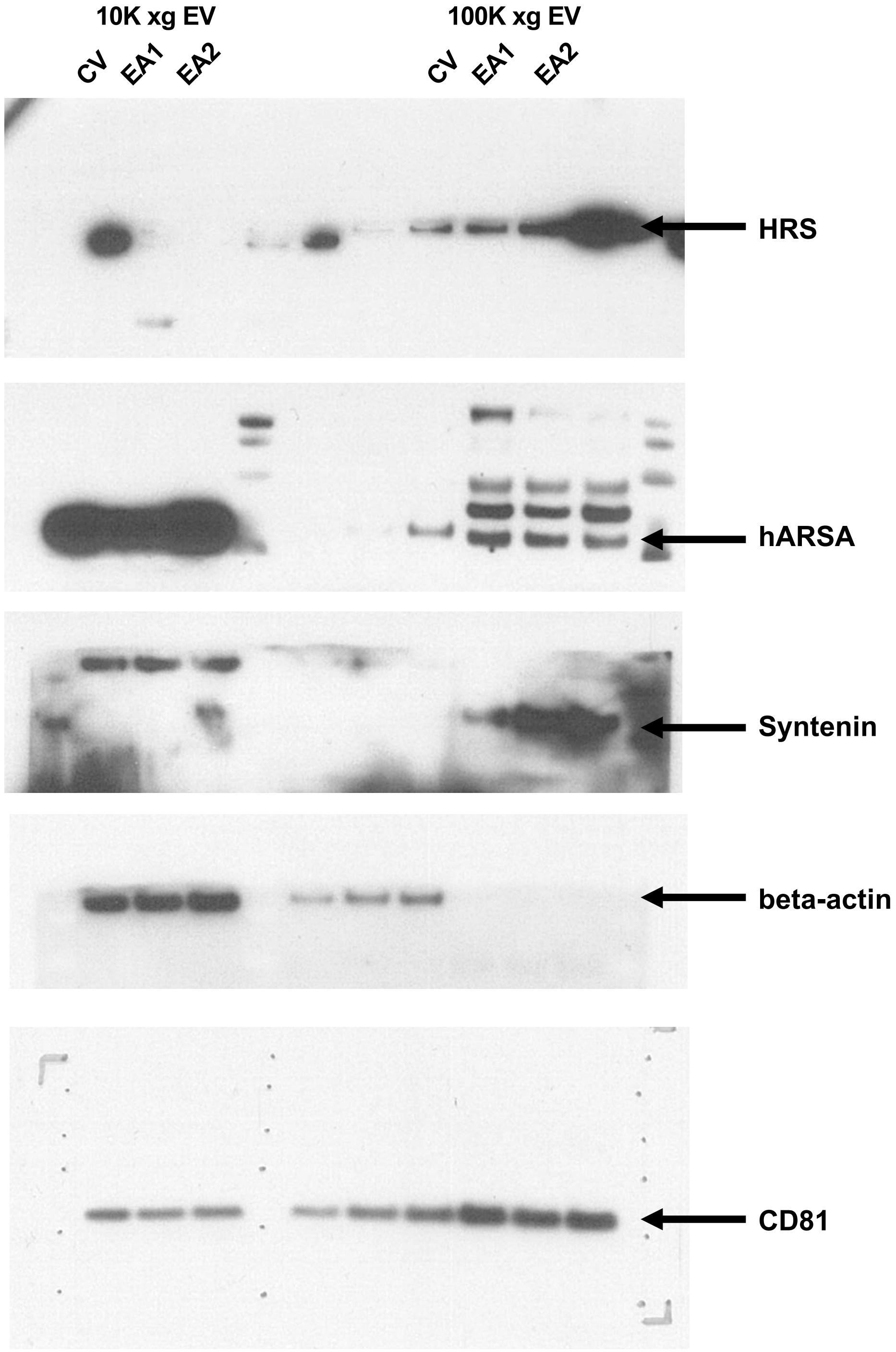

**Table S11**. Raw audioradiogram from supplemental Figure 5A.

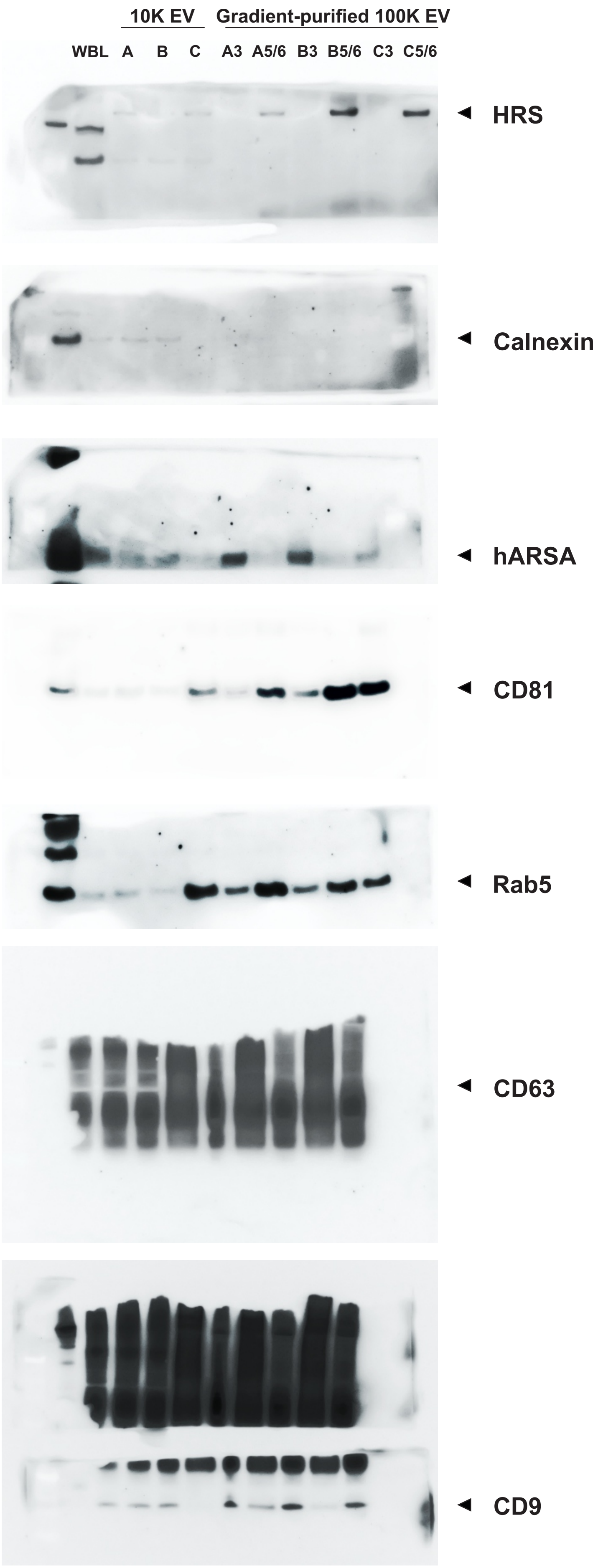

**Table S12**. Raw audioradiogram from supplemental Figure 5B.
